# Protein Language Model Identifies Disordered, Conserved Motifs Implicated in Phase Separation

**DOI:** 10.1101/2024.12.12.628175

**Authors:** Yumeng Zhang, Jared Zheng, Bin Zhang

**Affiliations:** Department of Chemistry, Massachusetts Institute of Technology, Cambridge, MA, USA; Department of Electrical Engineering and Computer Science, Massachusetts Institute of Technology, Cambridge, MA, USA

## Abstract

Intrinsically disordered regions (IDRs) play a critical role in phase separation and are essential for the formation of membraneless organelles (MLOs). Mutations within IDRs can disrupt their multivalent interaction networks, altering phase behavior and contributing to various diseases. Therefore, examining the evolutionary constraints of IDRs provides valuable insights into the relationship between protein sequences and phase separation. In this study, we utilized the ESM2 protein language model to map the residue-level mutational tolerance landscapes landscape of IDRs. Our findings reveal that IDRs, particularly those actively participating in phase separation, contain conserved amino acids. This conservation is evident through mutational constraints predicted by ESM2 and supported by direct analyses of multiple sequence alignments. These conserved, disordered amino acids include residues traditionally identified as “stickers” as well as “spacers” and frequently form continuous sequence motifs. The strong conservation, combined with their potential role in phase separation, suggests that these motifs may act as functional units under evolutionary selection to support stable MLO formation. Our findings underscore the insights into phase separation’s molecular grammar made possible through evolutionary analysis enabled by protein language models.

## Introduction

Membraneless organelles (MLOs), such as nucleoli, stress granules, and P-bodies, are distributed throughout diverse cellular environments and play a vital role in forming specialized biochemical compartments that drive essential cellular functions^1–13^ These biomolecular condensates typically assemble through phase separation, dynamically recruiting reactants and releasing products to improve the efficiency and specificity of cellular processes.^4,14,15^ Intrinsically disordered regions (IDRs), which lack well-defined tertiary structures yet exhibit unique structural disorder, frequently act as scaffolds within MLOs.^14,16–19^ IDRs facilitate multivalent interactions, including *π*-*π* stacking, cation-*π*, and electrostatic interactions, that promote phase separation. Mutations in IDRs can disrupt phase behavior, potentially leading to MLO dysfunction and contributing to diseases such as neurodegenerative disorders and cancer.^20–23^

Substantial research has focused on linking protein sequences to the phase behaviors of condensates.^4,6,7,9,17,24–42^ These studies support the “stickers and spacers” framework,^3,4,43–46^ in which specific residues, termed “stickers,” drive strong, specific interactions, while “spacer” regions act as flexible linkers with minimal nonspecific interactions.^37,47–50^ Sood and Zhang ^51^ further introduced an evolutionary dimension, proposing that IDRs adapt this framework to balance effective phase separation with compositional specificity. For instance, membrane-less organelles (MLOs) form through specific interactions among stickers, while minimizing non-specific interactions among spacers to enable condensate formation with defined structural and compositional properties. This evolutionary pressure supports the enrichment of low-complexity domains (LCDs) in IDRs, reducing nonspecific interactions yet preserving conserved stickers critical for condensate specificity and stability. This evolutionary perspective, while promising, has not been extensively validated.

Evolutionary analysis of IDRs is challenging due to difficulties in sequence alignment,^52–58^ though several studies have attempted alignment of disordered proteins with promising results.^59–61^ Recent advances in protein language models^62–67^ provide alternative approaches for sequence analysis and information decoding. Trained on large protein datasets, such as UniProt,^68^ these models leverage neural network architectures like the Transformer^69^ to capture correlations among amino acids within a sequence. These correlations enable the models to identify constraints, enforced by the whole sequence, that influence the chemical identity of amino acids at specific positions. Since these constraints often reflect functional or structural requirements, the mutational preferences predicted by these models can be interpreted as proxies for evolutionary constraint or mutational tolerance.^70–77^ Unlike traditional methods, these predictions do not rely on sequence homology, making them particularly attractive for analyzing disordered protein sequences where alignment is unreliable.

While protein language models have been widely applied to structured proteins,^73–75,77–80^ it is important to emphasize that these models themselves are not inherently biased toward folded domains. For example, the Evolutionary Scale Model (ESM2)^79^ is trained as a probabilistic language model on raw protein sequences, without incorporating any structural or functional annotations. Its unsupervised learning paradigm enables ESM2 to capture statistical patterns of residue usage and evolutionary constraints without relying on explicit structural information. Thus, the success of ESM2 in modeling the mutational landscapes of folded proteins^73–75,77–79^ reflects the model’s ability to learn sequence-level constraints imposed by natural selection — a property that is equally applicable to IDRs if those regions are also under functional selection. Indeed, protein language models are increasingly been used to analyze variant effects in IDRs.^70,71,78^

In this study, we employ the ESM2 model to investigate the mutational landscape of IDRs involved in MLO formation. Our analysis demonstrates the utility of ESM2 for examining disordered sequences, identifying a notable subset of amino acids that exhibit mutation resistance. These amino acids are evolutionarily conserved, as confirmed through multi-sequence alignment analysis. Notably, regions associated with phase separation are significantly enriched in these conserved residues. Importantly, the conserved disordered amino acids include both sticker residues, such as tyrosine (Y), tryptophan (W), and phenylalanine (F), and spacer residues, such as alanine (A), glycine (G), and Proline (P). These residues frequently colocalize within continuous sequence stretches, underscoring the functional relevance of entire motifs rather than isolated residues in MLO formation. Our findings provide strong evidence for evolutionary pressures acting on specific IDRs, likely to maintain their roles in scaffolding phase separation mechanisms.

## Results

### Protein Language Model for Quantifying the Mutational Landscape of MLO Proteins

To examine the mutational tolerance of IDRs and their connection to phase separation, we compiled a database of human proteins with disordered regions. From this, we identified a subset of 939 proteins associated with the formation of membraneless organelle, referred to as MLO-hProt. The Methods section provides additional information on the dataset preparation. The dataset contains proteins with varying numbers of disordered residues, ranging from a few dozen to several thousand per protein (Figure 1A). These proteins are involved in the assembly of various MLOs, including P-bodies, Cajal bodies, and centrosome granules, and are distributed across both nuclear and cytoplasmic compartments.

**Figure 1:**
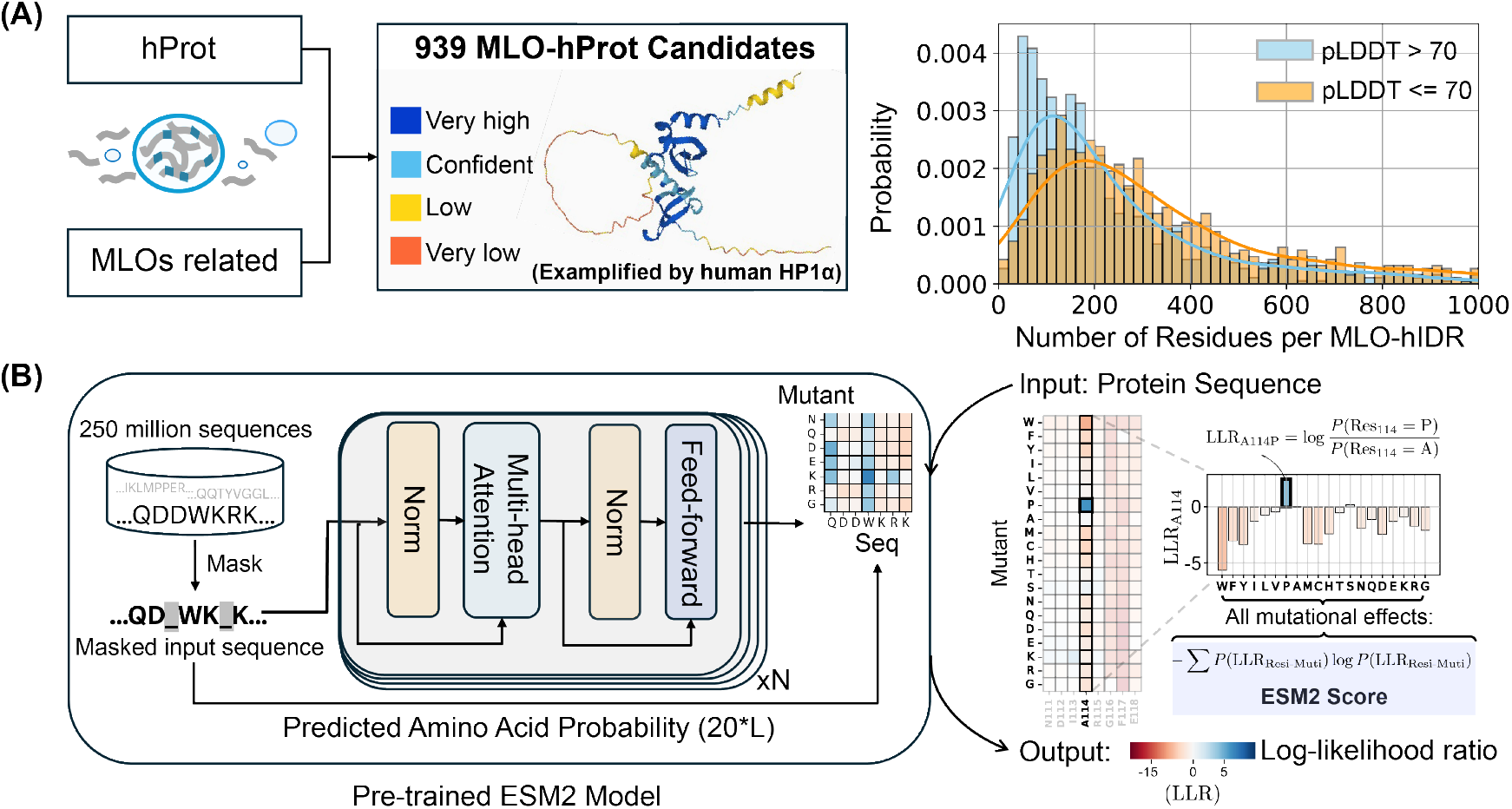
Predicting the mutational landscape of IDRs involved in MLO formation (MLO-hProt) using ESM2. (A) Schematic representation of the MLO-hProt database construction. The right panel shows the distribution of disordered and folded residues identified in each protein, where structural order is assigned using the AlphaFold2 pLDDT score.(B) Workflow of the pre-trained ESM2 model (left)^79^ for predicting the mutational landscape of a given protein sequence (right). Upon receiving a protein sequence as input, ESM2 generates a log-likelihood ratio (LLR) for each mutation type at each residue position. Using the 20-element LLR vector, we compute the ESM2 score for each residue (Eq. 1) to assess mutational tolerance.

We analyzed proteins in the MLO-hProt dataset using the protein language model, ESM2.^79^ As illustrated in Figure 1B, ESM2 is a conditional probabilistic model (masked language model) that predicts the likelihood of specific amino acids appearing at a given position, based on the surrounding sequence context.

The mutational tolerance of a specific amino acid at a given site is defined as follows. ESM2 enables the quantification of the probability, or likelihood, of observing any of the 20 amino acids at site *i*. To assess the preference for a mutant over the wild-type (WT) residue, we calculate the log-likelihood ratio (LLR) between the mutant and WT residues. Consequently, a 20-element vector representing the LLRs for each amino acid can be generated at each site (Figure 1B). This vector is then condensed into a single value, referred to as the ESM2 score, which is derived using an information entropy expression for the LLR probabilities of individual amino acids (Eq. 1 in the Methods Section).

The ESM2 score provides a measure of the overall mutational tolerance of a given residue. Lower scores indicate higher mutational constraint and reduced flexibility, implying that these residues are more likely essential for protein function, as they exhibit fewer permissible mutational states.

### ESM2 Identifies Conserved, Disorderd Residues

We next used ESM2 to analyze the mutational tolerance of amino acids in both structured and disordered regions. We carried out ESM2 predictions for all proteins in the MLO-hProt dataset, and determined the ESM2 scores of individual amino acids. In addition, to quantify structural disorder, we computed the AlphaFold2 predicted Local Distance Difference Test (pLDDT) scores for each residue. The pLDDT scores have been shown to correlate well with protein flexibility and disorder,^81^ making them a reliable tool for distinguishing structured from unstructured regions. Following previous studies,^82,83^ we used a threshold of pLDDT = 70 to differentiate ordered from disordered residues. This threshold reflects amino acid composition preferences for folded versus disordered proteins (see Figure S1).^82,83^

We first analyzed the relationship between ESM2 and pLDDT scores for human Heterochromatin Protein 1*α* (HP1*α*, residues 1–191). HP1*α* is a crucial chromatin organizer that promotes phase separation and facilitates the compaction of chromatin into transcriptionally inactive regions.^34,84–87^ HP1*α* comprises both structured and disordered segments, as illustrated in Figure 2A. Here, residues with pLDDT scores exceeding 70 (indicating ordered regions) are shown in white, while disordered segments (pLDDT ≤ 70) are highlighted in blue. Similar plots for other proteins are included in Figure S2. Figure 2B displays the AlphaFold2 predicted structure of HP1*α*, with residues colored according to their pLDDT scores.

**Figure 2:**
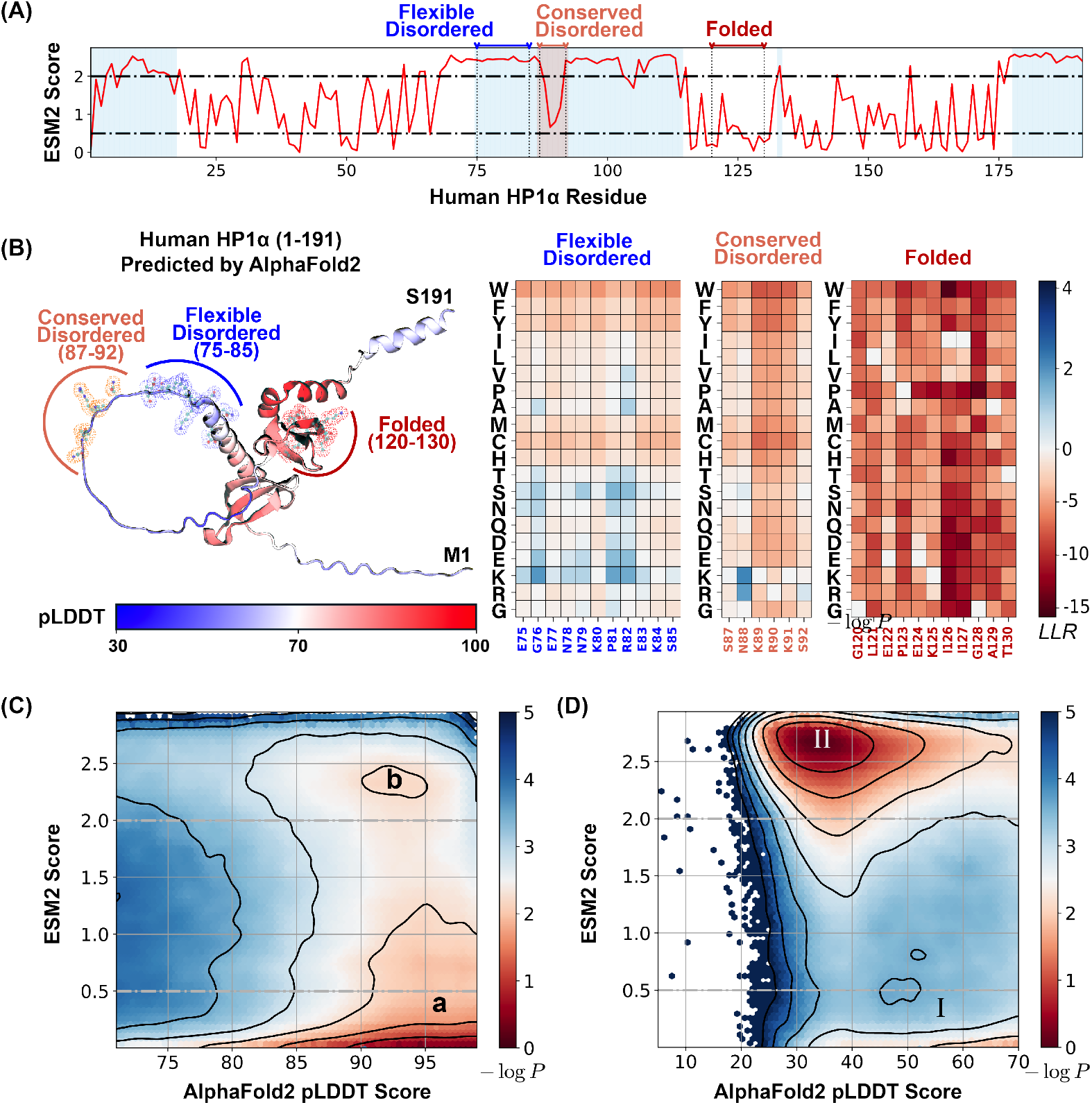
ESM2 predicts mutational landscape for structured and disordered residues. (A) The ESM2 scores for amino acids in the human HP1α protein (UniProt ID: P45973) are presented, with residues having pLDDT scores below 70 highlighted in blue to signify regions lacking a defined structure. (B) A detailed view of the mutational landscape across three regions with varying degrees of structural order. On the left, the AlphaFold2-predicted structure of the human HP1α protein is displayed in cartoon representation, with residues colored according to their pLDDT scores. Three specific regions, representing flexible disordered (residues 75–85), conserved disordered (residues 87–92), and folded (residues 120–130) segments, are highlighted in blue, orange, and red, respectively, using ball-and-stick styles. The panels on the right depict the ESM2 LLR predictions for each of these regions. (C, D) Histograms of pLDDT and ESM2 score distributions for structured (C) and disordered (D) residues are presented. Contour lines indicate free energy levelscomputed as*−* log *P* (pLDDT, ESM2), where *P* is the probability density of residues based on their pLDDT and ESM2 scores. Contours are spaced at 0.5-unit intervals to distinguish areas of differing density.

Our analysis demonstrated that HP1*α*’s structured domains consistently yield low ESM2 scores, reflecting strong mutational constraints characteristic of folded regions. These constraints are further evident in the local LLR predictions, as shown in Figure 2B, where we illustrate the folded region G120-T130. Given the critical role of maintaining the 3D conformation of structured domains, mutations with pronounced deleterious effects are likely to significantly disrupt protein folding. This interpretation aligns with prior studies that have reported a strong correlation between ESM2 LLRs and changes in free energy associated with protein structural stability.^78,79^

In contrast, disordered regions, including the N-terminal (residues 1–19), C-terminal (175–191), and hinge domain (70–117), typically exhibit higher ESM2 scores, indicating increased mutational flexibility. Nonetheless, not all disordered regions show similar flexibility. Within the hinge domain, a conserved segment known as the KRK patch (highlighted in orange)^84,88–90^ shows low ESM2 scores, despite being disordered. This distinction allows us to classify disordered regions into two types: “flexible disordered” regions, which show high ESM2 scores and greater mutational tolerance, and “conserved disordered” regions, which display low ESM2 scores, indicating varying levels of mutational constraint despite a lack of stable folding.

We then examined the distribution of ESM2 scores for all amino acids in the MLO-hProt dataset to evaluate the generality of these patterns. Amino acids in folded regions (pLDDT *>* 70) consistently yield low ESM2 scores, reflecting strong mutational constraints. As shown in Figure 2C, the histogram of ESM2 versus pLDDT scores for structured residues reveals a dominant population with low ESM2 values (region **a**, ESM2 Score *≤* 0.5), consistent with the established understanding that folded domains require structural and functional integrity and are thus more mutation-sensitive.^91–93^

In contrast, disordered residues (pLDDT *≤* 70) predominantly show high ESM2 scores (region **II**, ESM2 Score *≥* 2.0), consistent with the rapid evolution and higher mutational tolerance typical of disordered proteins.^94–96^ However, as shown in Figure 2D, a substantial subset of disordered amino acids also exhibit low ESM2 scores (region **I**). Given that low ESM2 scores generally reflect mutational constraint in folded proteins, the presence of region **I** among disordered residues suggests that certain disordered amino acids are evolutionarily conserved and likely functionally significant.

### ESM2 Scores Correlate with Sequence Conservation

Our analysis indicates that a substantial proportion of amino acids within disordered regions exhibit low mutational tolerance. To evaluate this hypothesis, we conducted an evolutionary analysis of MLO-hProt proteins, examining the conservation patterns of individual amino acids.

This analysis was based on a multi-sequence alignment (MSA) of homologs of MLO-hProt proteins. We employed HHblits for homolog detection, a method particularly suited to disordered proteins as it effectively captures distant sequence similarities in highly divergent sequences.^97–99^ The presence of folded domains in these proteins facilitates reliable alignment between references and their query homologs. To exclude sequences that no longer qualify as homologs, we filtered for sequences with at least 20% identity to the reference, resulting in homologous sets ranging from tens to thousands per protein (Figure 3A). From these aligned sequences, we calculated the conservation score for each reference amino acid as the ratio of its occurrence in homologs to the total number of sequences (see Eq. 2 in the Methods Section).

**Figure 3:**
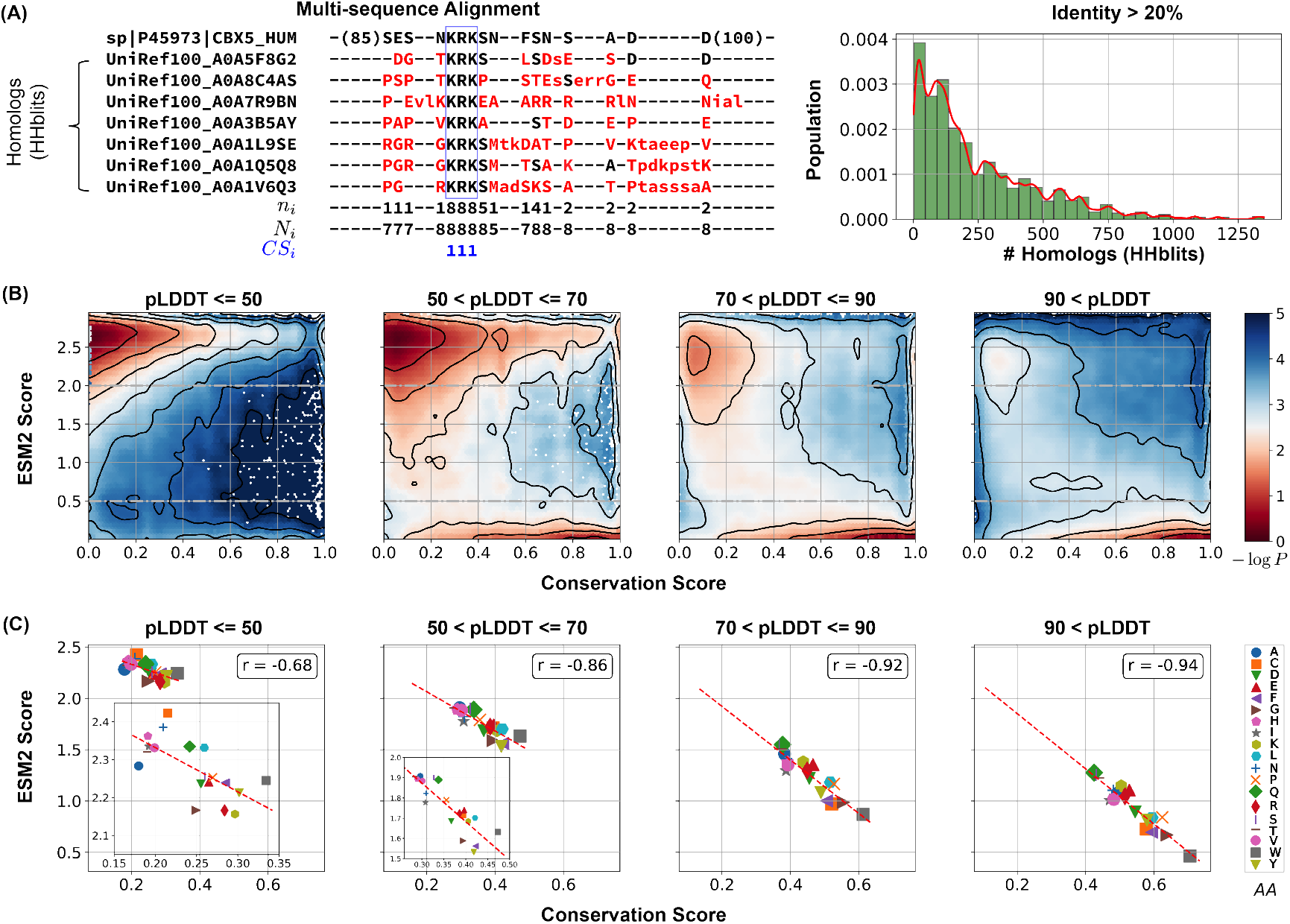
Low ESM2 scores correlate strongly with evolutionary conservation. (A) Estimating amino acid conservation using multiple sequence alignment. The conservation score calculation is demonstrated for human HP1α protein along with a subset of its homologs found by HHblits. In the aligned sequences, missing residues appear as dashed lines, insertions are shown in lowercase letters, and mismatches are highlighted in red. The three rows below the alignment indicate at position *i*: *n*_*i*_, the number of conserved residues from the reference sequence to the query sequences; *N*_*i*_; the total number of existing residues; and *CS*_*i*_, the conservation score calculated from Eq. 2, respectively. The right panel illustrates the distribution of homolog counts for each MLO-hProt found by HHblits. (B) Histograms showing conservation and ESM2 score distributions for all residues in MLO-hProt, grouped by pLDDT scores from AlphaFold2. The contour lines denote free energy levels, calculated as *−* log *P* (CS, ESM2), where *P* is the probability density of residues based on their conservation and ESM2 scores. Contours are spaced at 0.5-unit intervals to highlight regions of distinct density. (C) Correlation between mean conservation and ESM2 scores for amino acids classified by structural order levels. Pearson correlation coefficients, *r*, are reported in the legends.

Our findings reveal a strong correlation between ESM2 scores and conservation scores. In Figure 3B, we present the histograms of ESM2 and conservation scores for all amino acids from MLO-hProt proteins. Given that folded domains generally show higher conservation scores than disordered regions, we further classified residues into four groups based on their AlphaFold2 pLDDT scores to assess conservation patterns across varying levels of structural disorder. This stratification allowed us to analyze conservation trends in detail. Across all categories, we observed bimodal distributions, reinforcing the correlation between increasing ESM2 scores and decreasing conservation.

The conservation of amino acids with low ESM2 scores is also apparent in Figure 3C. For each of the four structural order groups, we computed the average ESM2 and conservation scores for all 20 amino acid types. Methionine (M) was excluded from the correlation analysis due to its frequent position as the initial residue in sequences, which complicates reliable mutational effect predictions.^70,100^ In each group, we consistently observed a strong correlation between average ESM2 and conservation scores.

While ESM2 scores align closely with conservation scores, the relative conservation of specific amino acids varies across structural order groups. In more disordered regions, hydrophilic residues such as Glutamine (Q), Lysine (K), and Arginine (R) exhibit lower ESM2 scores, indicating that mutations in these residues are particularly detrimental. Conversely, hydrophobic residues like Valine (V) and Isoleucine (I) show higher ESM2 scores, suggesting they experience reduced evolutionary constraints. In more folded domains, hydrophobic residues such as W and F are more conserved (see Figure S3), consistent with the characteristic conservation patterns of proteins across different disorder levels.^101,102^ Overall, these findings strongly support our hypothesis that ESM2 scores effectively capture evolutionary conservation, enabling the identification of functionally significant residues through the mutational landscape, independent of structural flexibility.

### Regions Driving Phase Separation Are Enriched with Conserved, Disordered Residues

The presence of evolutionarily conserved disordered residues raises the question of their functional significance. To explore this, we identified disordered regions of MLO-hProt using a pLDDT score ≤ 70 and partitioned these regions into two categories: drivers (dMLO-hIDR), which actively drive phase separation, and clients (cMLO-hIDR), which are present in MLOs under certain conditions but do not promote phase separation themselves.^103^ Additionally, IDRs from human proteins not associated with MLOs, termed nMLO-hIDR, were included as a control. To enhance statistical robustness, we extended our dataset by incorporating driver proteins from additional species,^104^ resulting in the expanded dMLO-IDR dataset. Figure S4 shows the amino acid composition across these datasets. Beyond the pLDDT-based classification, the majority of residues in these datasets are also predicted to be disordered by various computational tools and supported by experimental evidence (Figures S5 and S6).

As illustrated in Figure 4A, there is a progressive increase in the fraction of conserved disordered residues and a corresponding decline in flexible disordered residues from non-phase-separating proteins (nMOL-hIDR) to clients (cMLO-hIDR) and drivers (dMLO-hIDR and dMLO-IDR) (see also Figure S7). Driver proteins, particularly those in the expanded dataset, display a notable reduction in flexible residues. These findings imply that disordered regions with a role in phase separation tend to contain functionally significant and evolutionarily conserved regions.

**Figure 4:**
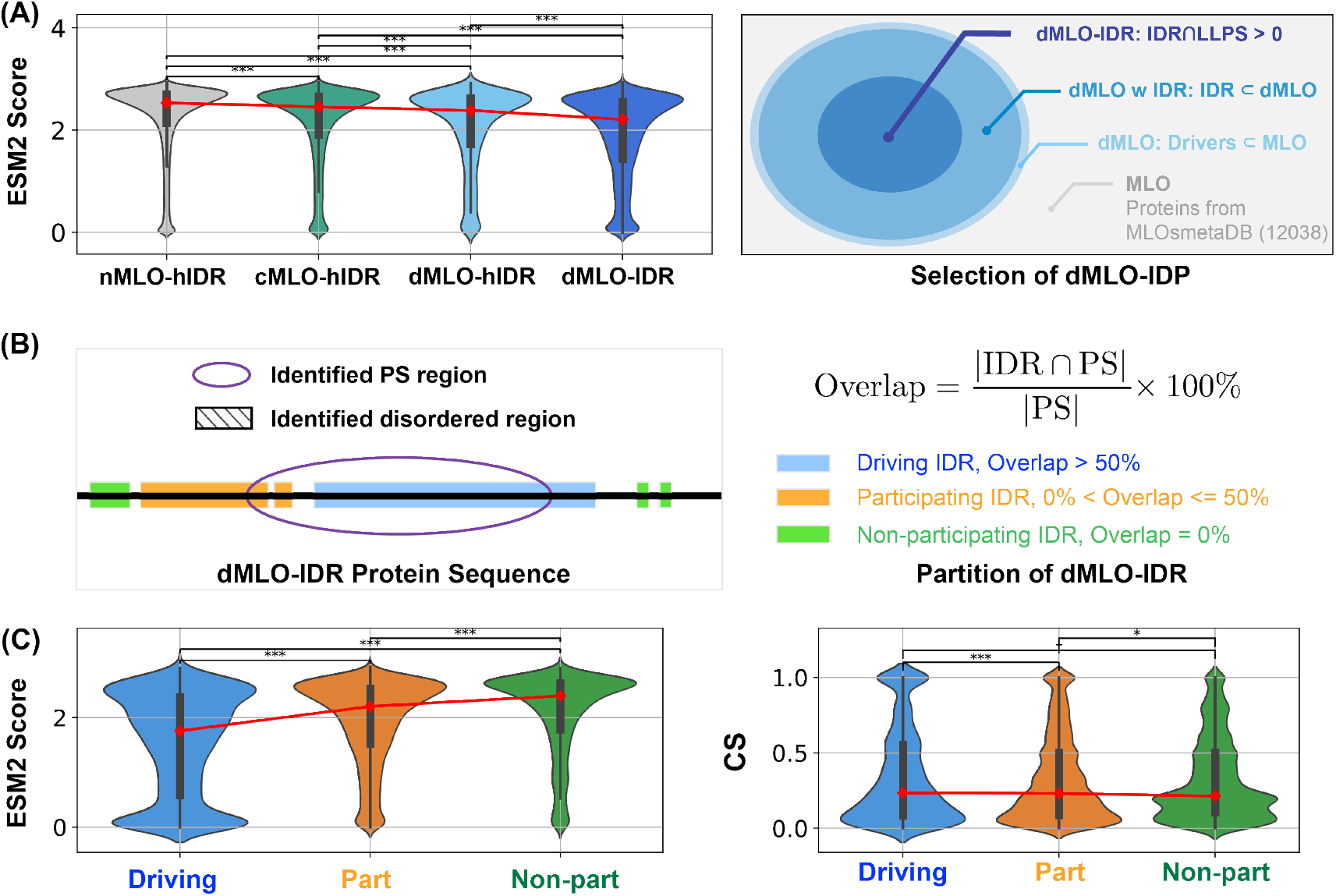
Phase separation driving IDRs exhibit more conserved disorder. (A) Population of ESM2 score for disordered residues in proteins from nMLO-hIDR, cMLO-hIDR, dMLO-hIDR, and dMLO-IDR datasets. Red dots indicate the mean values of the respective distributions. The selection of proteins in the dMLO-IDR dataset is shown in the right panel. See also methods for details in dataset preparation. (B) The classification of three IDR functional groups based on their overlap with the experimentally identified phase separation (PS) segments. (C) The distribution of the ESM2 score for residues in three IDR groups, driving (blue), participating (orange), and non-participating (green) shown in the violin plot. (D) The distribution of the conservation score (CS) for residues in three IDR groups shown in the violin plot with same coloring scheme in C. Pairwise statistical comparisons were conducted using two-sided Mann–Whitney U tests on the ESM2 score distributions (null hypothesis: the two groups have equal medians). P-values indicate the probability of observing the observed rank differences under the null hypothesis. Statistical significance is denoted as follows: **\*\*\***: p < 0.001; **\*\***: p < 0.01; **\***:p < 0.05; **†**: p < 0.10 (marginal); **n.s**.: (not significant, p *≥* 0.10).

We further examined the sequence location of conserved, disordered residues in driver proteins (dMLO-IDR). For these proteins, experimentally verified segments have been identified, the deletion or mutation of which impairs phase separation^104–108^ (Figure 4B). These segments can include both structured and disordered regions. Herein, if a disordered region constitutes over 50% of the phase-separating segment, we designate it as “driving”, indicating a likely critical contribution to phase separation. If the disordered region represents less than 50%, we classify it as “participating”, with a potentially limited role. Finally, if there is no overlap between the disordered region and the phase-separating segment, we categorize it as “non-participating”. The number of segments in the three IDR groups, along with their amino acid compositions, are shown in Figure S8.

We then analyzed the distribution of ESM2 predictions across these IDR groups. In alignment with Figure 3A, we observed a significantly higher proportion of conserved disordered residues within driving IDRs, while few were present in non-participating IDRs. Supporting the ESM2 predictions, conservation analysis based on MSA also indicated that drving IDRs contain a greater concentration of conserved residues (Figure 4C). Collectively, these findings demonstrate that ESM2 effectively identifies evolutionarily conserved functional sites, enriched in IDR regions likely involved in driving phase separation.

### Conserved, Disordered Residues Form Motifs

Finally, we investigated the chemical identities of conserved residues within driving IDRs to understand their potential role in phase separation. Figure 5A displays the average ESM2 LLR predictions for each of the 20 amino acids in the mutational matrix, indicating that mutations to most amino acids are generally unfavorable, as reflected by their low, negative LLR values. This trend is particularly pronounced in driving IDRs compared to nMLO-IDRs or non-participating IDRs (Figure S9).

**Figure 5:**
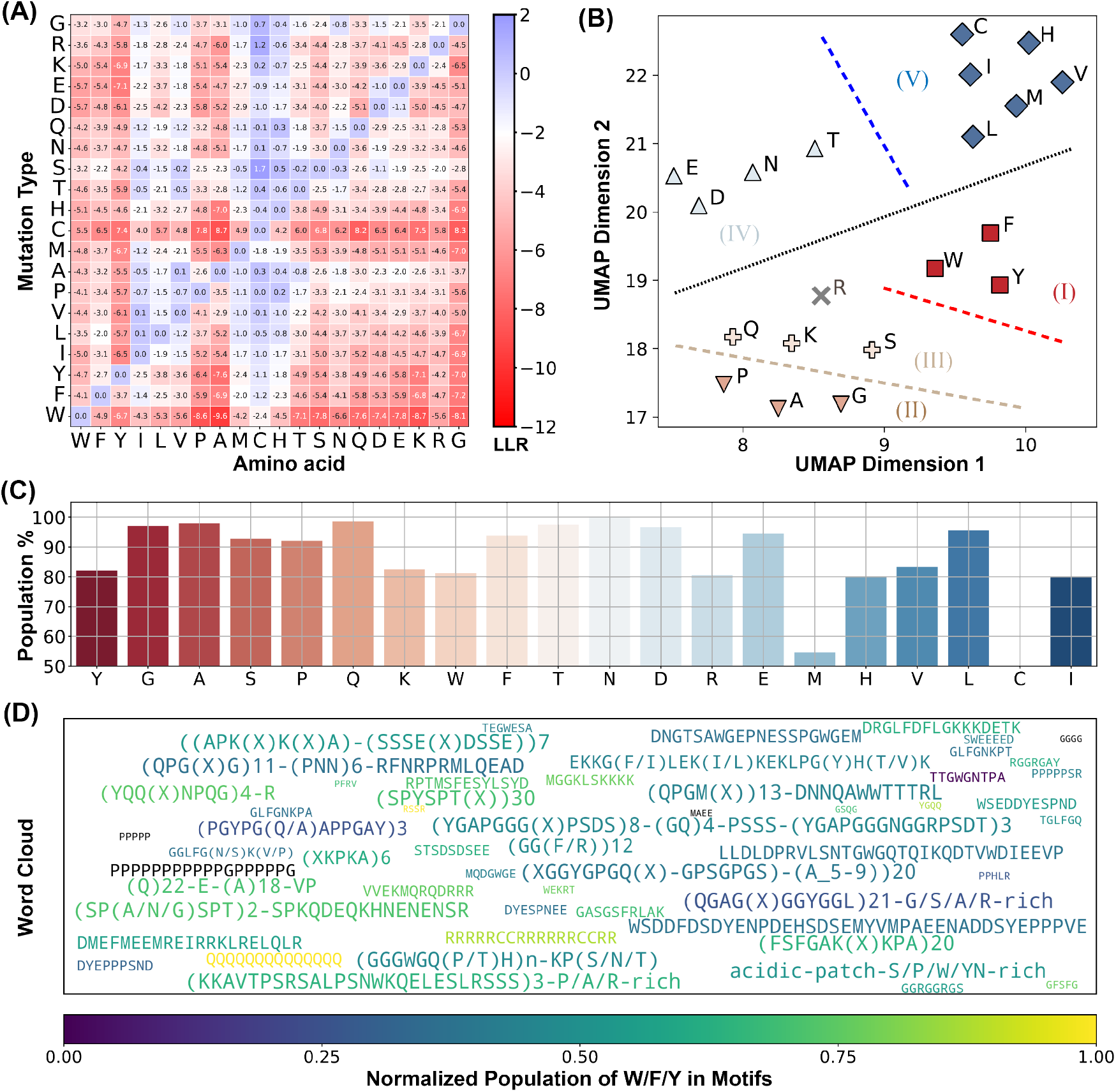
ESM2 Identifies conserved Motifs in driving IDRs. (A) Mean LLR values for the 20 amino acids calculated by averaging across all residues of each amino acid type. (B) Clustering of amino acids based on the two UMAP embeddings of their LLR vectors presented in part A. The dashed lines are manually added for a clear visualization of the separation of each group. (C) The percent of conserved residues locating in motifs for all amino acids. (D) Word cloud of motifs identified by ESM2. The word font size reflects the relative motif length, while the color represents the proportion of “sticker” residues (Y, F, W, R, K, and Q) within each motif.

We further characterize these conserved residues within driving IDRs. Using hierarchical clustering on two UMAP-derived embeddings from the LLR vectors, we grouped amino acids into five clusters (Figure 5B). This approach distinguishes more conserved residues (Groups I to III) from the more flexible residues (Groups IV and V). Notably, W, F, and Y—often referred to as “stickers” due to their crucial role in phase separation^43,109–111^—are uniquely grouped within the highly conserved Group I. These findings support the expectation that amino acids essential to phase separation are often evolutionarily conserved, aligning with their central role in functional stability.

Interestingly, residues in Groups II and III, which include traditional “spacers” (G, A, P, and S), also show high conservation and resistance to most mutation types, particularly hydrophobic mutations (Figure 5A). Spacer residues, generally regarded as less critical for interactions driving condensate formation, were unexpectedly conserved, suggesting a broader functional relevance than previously assumed.

We propose that this conservation pattern for spacers is likely not due to isolated residue conservation but may instead reflect the conservation of specific sequence stretches. To examine this hypothesis, we identified ESM2 “motifs” as continuous sequence regions with average ESM2 scores below 0.5. A full list of motifs is available in the appendix (`ESM2_motif_with_exp_ref.csv`). We observed that conserved amino acids with ESM2 scores below 0.5 are predominantly located within these motifs (see Figure 5C and Figure S10A). For instance, conserved glycine residues have a 97.9% likelihood of being part of an ESM2 motif, with similar probabilities observed for other spacers, such as alanine (99.0%) and proline (93.1%), as well as for sticker residues like Y, W, and F.

These results suggest that IDRs crucial for phase separation frequently contain conserved sequence motifs composed of both sticker and spacer residues. Interestingly, many of these motifs have been experimentally validated as essential for phase separation, with representative motifs for each driving IDR listed in `ESM2_motif_with_exp_ref.csv`. In these cases, mutations or deletions have been shown to disrupt phase separation. For visualization, a word cloud of these motifs is presented in Figure 5D. Altogether, our analysis suggest a tendency for IDRs to uniquely cluster conserved residues into motifs and execute significant biological roles in phase separation.

## Conclusions and Discussion

We have utilized the protein language model ESM2 to investigate the mutational landscape of IDRs. Our analysis reveals a substantial population of mutation-resistant amino acids. Multi-sequence alignment confirms their evolutionary conservation. Notably, regions actively involved in phase separation are enriched with these conserved, disordered residues, supporting their potential role in the formation of membraneless organelles. These findings underscore evolutionary constraints on specific IDRs to preserve their functional roles in scaffolding phase separation processes.

We emphasize that the results presented in Figure 4 do not directly demonstrate that conserved residues are preferentially located in regions associated with phase separation. Although these regions are more enriched in conserved amino acids, their total sequence length can be smaller than that of non-phase-separating regions. As a result, the absolute number of conserved residues may still be higher outside phase-separating regions. To quantitatively assess this, we calculated, for each protein in the MLO-hProt dataset, the probability *p* of finding conserved residues within regions contributing to phase separation. These regions include both “driving” and “participating” segments, as defined in Figure 4 of the main text. Figure S11 shows the distribution of *p* across all proteins. For comparison, we also present the distribution of 1 *− p*, which reflects the probability of finding conserved residues in non-phase-separating regions. While the average value of *p* exceeds 0.5, indicating a trend toward conserved residues being more frequently located in phase-separating regions, the difference between the two distributions is not statistically significant. Future studies with expanded datasets may be necessary to clarify this trend.

### Related Work

Numerous studies have sought to identify functionally relevant amino acid groups within IDRs.^11,34,43,46,109,112–116^ For instance, using multiple sequence alignment, several groups have identified evolutionarily conserved residues that contribute to phase separation.^13,59,61,84,113,117–121^ Alderson et al. employed AlphaFold2 to detect disordered regions with a propensity to adopt structured conformations, suggesting potential functional relevance.^83^

In contrast, our approach based on ESM2 is more direct: it identifies conserved residues without relying on alignment or presupposing that functional significance requires folding into stable 3D structures. Notably, many of the conserved residues identified in our analysis exhibit low pLDDT scores (Figure 2), implying potential functional roles independent of stable conformations.

### Relationship to the Stickers and Spacers Model

To interpret the functional implications of conserved residues, we contextualize our findings within the stickers-and-spacers framework. Our results indicate that these conserved amino acids encompass both “sticker” and “spacer” classifications, as defined in recent literature.^13,84,111,113,117–119,121–123^ This suggests that residue-level classification may obscure functional roles in MLO formation.

Instead, evolutionarily conserved motifs, which can be readily identified through ESM2 score profiles, likely represent functionally significant units that integrate combinations of stickers and spacers. Perturbing these motifs in in vivo systems may offer a productive strategy for elucidating their biological functions.

Outside the conserved motifs, IDRs display greater mutational tolerance, with a general preference for hydrophilic residues. This structural arrangement—functionally constrained motifs embedded in more flexible, mutation-tolerant regions—is consistent with a generalized stickers-and-spacers framework. Conserved motifs likely engage in stronger interactions that drive phase separation, while interactions among non-motif regions remain attenuated. This spatial organization supports the formation of MLOs with defined chemical compositions and interaction specificity, as proposed by Sood and Zhang ^51^.

### Future Directions

Several promising directions could extend this work, both to refine our mechanistic understanding and to explore clinical relevance. One avenue is testing the hypothesis that conserved motifs in scaffold proteins act as functional stickers, mediating strong intermolecular interactions. This could be evaluated computationally via free energy calculations or experimentally via interaction assays. Deletion of such motifs in client proteins may also reduce their partitioning into condensates, illuminating their roles in molecular recruitment.

To explore potential clinical implications, we analyzed pathogenicity data from ClinVar.^124,125^ As shown in Figure S12A, single-point mutations with low LLR values—indicative of constrained residues—are enriched among clinically reported pathogenic variants, while benign variants typically exhibit higher LLR values. Moreover, mutations within conserved motifs are significantly more likely to be pathogenic than those in non-motif regions (Figure S12B). These findings highlight the potential of ESM2 as a first-pass screening tool for identifying clinically relevant residues and suggest that the conserved motifs described here may serve as priorities for future studies, both mechanistic and therapeutic.

### Limitations

Despite these promising findings, our study has several limitations. Most notably, the analysis is entirely computational, relying on ESM2-derived predictions and sequence-based conservation without direct experimental validation. While the strong correlation between ESM2 scores and evolutionary conservation suggests functional constraint, the specific biological roles of many conserved motifs remain uncharacterized.

The motifs we identify are located within broader regions that have previously been shown, through experimental studies, to drive phase separation. This provides circumstantial support for their functional relevance. However, these motifs typically represent only a small fraction of the larger disordered segments implicated in phase separation. As such, it is not possible to attribute the functional effect of the entire region solely to the identified motifs—other segments of the sequence may also make essential contributions. Therefore, while suggestive, the current evidence does not constitute direct experimental validation of motif-level functionality.

Future studies should aim to dissect the roles of these motifs in isolation and in context, using targeted mutagenesis, biophysical assays, and cellular perturbations. Such work will be crucial for establishing whether the conserved motifs identified here act as discrete functional units or as components of a broader sequence context required for biomolecular condensation and related processes.

## Methods

### Data Collection and Preprocessing

#### Human protein dataset

To construct the MLO-hProt dataset, we initially identified human proteins with disordered residues using the UniProtKB database.^126^ From these proteins, we selected candidates with at least 10% of their residues exhibiting a pLDDT score ≤ 70, yielding a total of 5,121 candidates. These were then cross-referenced with entries in the CD-code Database,^127^ yielding a final subset of 939 proteins associated with membraneless organelle formation (Figure 1A). The remaining 4,182 proteins, which were not linked to membraneless organelle formation, comprise the nMLO-hProt dataset. We further identified disordered regions from the datasets by selecting residues with pLDDT score ≤ 70, refereed as MLO-hIDR and nMLO-hIDR. The CD-code Database also classifies the MLO-hProt proteins into two categories: drivers (n = 82) and clients (n = 814), with the corresponding disordered regions labeled as dMLO-hIDR and cMLO-hIDR. Additional information on MLO-hIDR candidate proteins and their biological roles is available in the appendix file `MLO-hIDR.csv`.

#### dMLO-IDR dataset

In addition to datasets comprising only human proteins, we developed a specialized dMLO-IDR database incorporating proteins from diverse species involved in driving phase separation. This database includes all Driver proteins cataloged in the MLOsMetaDB database,^104^ which documents experimentally validated disordered and phase-separating regions. Beginning with 780 Driver proteins, we filtered the dataset to retain entries where disordered regions overlap with phase-separating segments, yielding 399 candidates across 40 species (Figure 4A). To further refine the dataset, we analyzed sequence identity, excluding homologous pairs with sequence identity exceeding 50%. This process resulted in a final set of 341 non-redundant candidates. These proteins play critical roles in mediating the formation of various MLOs, including P-bodies, stress granules, paraspeckles, and centrosomes (see Appendix: `dMLO-IDR.csv`).

### AlphaFold2 score for structural order

The pLDDT scores for human proteins were retrieved from the AlphaFold Protein Structure Database^128^ by selecting the *Homo sapiens* organisms (Reference Proteome ID: UP000005640).

### ESM2 predictions for Mutational Preferences

We employed the code and pretrained parameters for ESM2 available from the model’s official GitHub repository at https://github.com/facebookresearch/esm to conduct mutational predictions. To optimize computational efficiency, we utilized the `esm2_t33_650M_UR50D` model, which has 650 million parameters and achieves prediction accuracy comparable to the larger 15B parameter model.^78^ In addition to calculating the log-likelihood ratio (LLR) for individual mutations, we defined an ESM2 score at each position to quantify mutational tolerance, formulated as

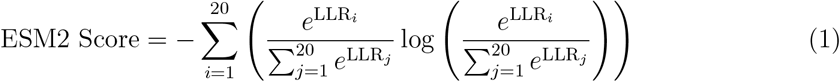

where LLR_*i*_ denotes the LLR value for the *i*-th amino acid mutation type.

### Evolutionary Sequence Analysis

We performed multi-sequence alignment (MSA) analysis using `HHblits` from the HH-suite3 software suite,^129–132^ a widely used open-source toolkit known for its sensitivity in detecting sequence similarities and identifying protein folds. HHblits builds MSAs through iterative database searches, sequentially incorporating matched sequences into the query MSA with each iteration. Sequence alignment was performed using the full-length protein sequences, encompassing both folded and disordered regions.

The HH-suite3 software was obtained from its GitHub repository (https://github.com/soedinglab/hh-suite). Homologous sequences were identified through searches against the UniRef30 protein database (release 2023/02).^133^ For each query, we performed three iterations of HHblits searches, incorporating sequences from profile HMM matches with an E-value threshold of 0.001 into the query MSA in each cycle. Using a lower E-value threshold (closer to 0) ensures greater sequence similarity among the matches, while multiple iterations enhance the alignment’s depth and accuracy. The resulting alignments in A3M format were converted to CLUSTAL format using the `reformat.pl` script provided in HH-suite, aligning all sequences to a uniform length (Figure 3A).

To refine alignment quality by focusing on closely related homologs, we filtered out sequences with *≤* 20% identity to the query, excluding weakly related sequences where only short segments show similarity to the reference. For each sequence, we calculated the percent identity by counting the number of positions, denoted as *n*, at which the amino acid matches the reference. The percent identity was then computed as *n/N*, where *N* represents the total length of the aligned reference sequence. This total includes residues in folded and disordered regions, as well as gap positions introduced during alignment.

The conservation score CS_*i*_ for position *i* was then calculated from the MSA as

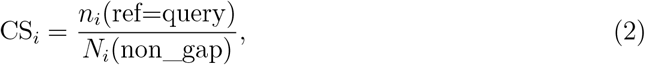

where *n*_*i*_(ref=query) represents the number of times the residues from the reference sequence appear across all sequences, and *N*_*i*_(non_gap) represents the total non-gap residues across the aligned sequences.

### Motif Identification

We defined motifs as contiguous stretches of amino acid sequences with an average ESM2 score of 0.5 or lower. To identify motifs within a given IDR, we implemented the following iterative procedure. Starting from either the N– or C–terminus of the sequence, we first locate the initial residue *i* whose ESM2 score falls within 0.5. From *i*, residues are sequentially appended in the direction toward the opposite terminus until the segment’s average ESM2 score exceeds 0.5; the first residue to breach this threshold is denoted *j*. The segment (*i, i* + 1, …, *j* − 1) is then recorded as a candidate motif. This process repeats starting from *j* until the end of the IDR is reached.

We perform this full procedure independently from both termini and designate the final motif as the intersection of the two candidate-motif sets. This bidirectional overlap strategy excludes terminal residues that might transiently satisfy the average-score criterion only due to adjacent low-scoring regions, thereby isolating the conserved core of each motif. All other residues—those not included in either directional pass—are classified as non-motif regions, minimizing peripheral artifacts.

When two motifs are in close proximity along the sequence, they may be merged into a single motif. Specifically, if the starting position of one motif is within 8 residues of the ending position of another, we define a candidate segment as the sequence spanning both motifs and the intervening residues. If the candidate segment’s average ESM2 score is below 0.5, it is included as a merged motif, replacing the individual motifs in the final list (Appendix: `ESM2_motif_with_exp_ref.csv`). In the analyses shown in Figure 5, we showed all motifs with *n* ≥ 4; however, varying motif minimal length *n* does not alter the overall conclusions (Figure S10B).

## Acknowledgement

This work was supported by the National Institutes of Health (Grant R35GM133580).

## Data and materials availability

https://github.com/ZhangGroup-MITChemistry/ESM2_IDR_LLPS

## References

(1) Hirose, T.; Ninomiya, K.; Nakagawa, S.; Yamazaki, T. A Guide to Membraneless Organelles and Their Various Roles in Gene Regulation. Nature Reviews Molecular Cell Biology 2023, 24, 288–304.

(2) Banani, S. F.; Lee, H. O.; Hyman, A. A.; Rosen, M. K. Biomolecular Condensates: Organizers of Cellular Biochemistry. Nature Reviews Molecular Cell Biology 2017, 18, 285–298.

(3) Choi, J.-M.; Holehouse, A. S.; Pappu, R. V. Physical Principles Underlying the Complex Biology of Intracellular Phase Transitions. Annual Review of Biophysics 2020, 49, 107–133.

(4) Pappu, R. V.; Cohen, S. R.; Dar, F.; Farag, M.; Kar, M. Phase Transitions of Associative Biomacromolecules. Chemical Reviews 2023, 123, 8945–8987.

(5) Ginell, G. M.; Holehouse, A. S. In Phase-Separated Biomolecular Condensates: Methods and Protocols; Zhou, H.-X., Spille, J.-H., Banerjee, P. R., Eds.; Springer US: New York, NY, 2023; pp 95–116.

(6) Latham, A. P.; Zhang, B. Maximum Entropy Optimized Force Field for Intrinsically Disordered Proteins. Journal of Chemical Theory and Computation 2020, 16, 773–781.

(7) Latham, A. P.; Zhang, B. Molecular Determinants for the Layering and Coarsening of Biological Condensates. Aggregate 2022, 3, e306.

(8) Latham, A. P.; Zhang, B. Unified Protein Force Field for Simulations of Liquid-Liquid Phase Separation. Biophysical Journal 2022, 121, 356a–357a.

(9) Latham, A. P.; Zhang, B. On the Stability and Layered Organization of Protein-DNA Condensates. Biophysical Journal 2022, 121, 1727–1737.

(10) Latham, A. P.; Zhu, L.; Sharon, D. A.; Ye, S.; Willard, A. P.; Zhang, X.; Zhang, B. Microphase Separation Produces Interfacial Environment within Diblock Biomolecular Condensates. eLife 2024, 12.

(11) Liu, S.; Wang, C.; Zhang, B. Toward Predictive Coarse-Grained Simulations of Biomolecular Condensates. Biochemistry 2025, 64, 1750–1761.

(12) Lao, Z.; Zhang, B. OpenNucleome for High-Resolution Nuclear Structural and Dynamical Modeling. Biophysical Journal 2024, 123, 297a.

(13) Jiang, H.; Wang, S.; Huang, Y.; He, X.; Cui, H.; Zhu, X.; Zheng, Y. Phase Transition of Spindle-Associated Protein Regulate Spindle Apparatus Assembly. Cell 2015, 163, 108–122.

(14) Hyman, A. A.; Weber, C. A.; Jülicher, F. Liquid-Liquid Phase Separation in Biology. Annual Review of Cell and Developmental Biology 2014, 30, 39–58.

(15) Feric, M.; Vaidya, N.; Harmon, T. S.; Mitrea, D. M.; Zhu, L.; Richardson, T. M.; Kriwacki, R. W.; Pappu, R. V.; Brangwynne, C. P. Coexisting Liquid Phases Underlie Nucleolar Subcompartments. Cell 2016, 165, 1686–1697.

(16) Borcherds, W.; Bremer, A.; Borgia, M. B.; Mittag, T. How Do Intrinsically Disordered Protein Regions Encode a Driving Force for Liquid–Liquid Phase Separation? Current Opinion in Structural Biology 2021, 67, 41–50.

(17) Mittag, T.; Pappu, R. V. A Conceptual Framework for Understanding Phase Separation and Addressing Open Questions and Challenges. Molecular Cell 2022, 82, 2201–2214.

(18) Dignon, G. L.; Best, R. B.; Mittal, J. Biomolecular Phase Separation: From Molecular Driving Forces to Macroscopic Properties. Annual Review of Physical Chemistry 2020, 71, turoverov.

(19) Turoverov, K. K.; Kuznetsova, I. M.; Fonin, A. V.; Darling, A. L.; Zaslavsky, B. Y.; Uversky, V. N. Stochasticity of Biological Soft Matter: Emerging Concepts in Intrinsically Disordered Proteins and Biological Phase Separation. Trends in Biochemical Sciences 2019, 44, 716–728.

(20) Alberti, S.; Dormann, D. Liquid–Liquid Phase Separation in Disease. Annual Review of Genetics 2019, 53, 171–194.

(21) Zbinden, A.; Pérez-Berlanga, M.; De Rossi, P.; Polymenidou, M. Phase Separation and Neurodegenerative Diseases: A Disturbance in the Force. Developmental Cell 2020, 55, 45–68.

(22) Rai, S. K.; Savastano, A.; Singh, P.; Mukhopadhyay, S.; Zweckstetter, M. Liquid-Liquid Phase Separation of Tau: From Molecular Biophysics to Physiology and Disease. Protein Science: A Publication of the Protein Society 2021, 30, 1294–1314.

(23) Das, S.; Lin, Y.-H.; Vernon, R. M.; Forman-Kay, J. D.; Chan, H. S. Comparative Roles of Charge, π, and Hydrophobic Interactions in Sequence-Dependent Phase Separation of Intrinsically Disordered Proteins. Proceedings of the National Academy of Sciences 2020, 117, 28795–28805.

(24) Brangwynne, C. P.; Tompa, P.; Pappu, R. V. Polymer Physics of Intracellular Phase Transitions. Nature Physics 2015, 11, 899–904.

(25) Zeng, X.; Pappu, R. V. Developments in Describing Equilibrium Phase Transitions of Multivalent Associative Macromolecules. Current Opinion in Structural Biology 2023, 79, 102540.

(26) Schmit, J. D.; Bouchard, J. J.; Martin, E. W.; Mittag, T. Protein Network Structure Enables Switching between Liquid and Gel States. Journal of the American Chemical Society 2020, 142, 874–883.

(27) Dzuricky, M.; Roberts, S.; Chilkoti, A. Convergence of Artificial Protein Polymers and Intrinsically Disordered Proteins. Biochemistry 2018, 57, 2405–2414.

(28) Flory, P. J. Thermodynamics of High Polymer Solutions. The Journal of Chemical Physics 1942, 10, 51–61.

(29) Huggins, M. L. Some Properties of Solutions of Long-chain Compounds. The Journal of Physical Chemistry 1942, 46, 151–158.

(30) Dignon, G. L.; Zheng, W.; Kim, Y. C.; Best, R. B.; Mittal, J. Sequence Determinants of Protein Phase Behavior from a Coarse-Grained Model. PLOS Computational Biology 2018, 14, e1005941.

(31) Dignon, G. L.; Zheng, W.; Mittal, J. Simulation Methods for Liquid–Liquid Phase Separation of Disordered Proteins. Current Opinion in Chemical Engineering 2019, 23, 92–98.

(32) Joseph, J. A.; Reinhardt, A.; Aguirre, A.; Chew, P. Y.; Russell, K. O.; Espinosa, J. R.; Garaizar, A.; Collepardo-Guevara, R. Physics-Driven Coarse-Grained Model for Biomolecular Phase Separation with near-Quantitative Accuracy. Nature computational science 2021, 1, 732–743.

(33) Regy, R. M.; Thompson, J.; Kim, Y. C.; Mittal, J. Improved Coarse-Grained Model for Studying Sequence Dependent Phase Separation of Disordered Proteins. Protein Science 2021, 30, 1371–1379.

(34) Latham, A. P.; Zhang, B. Consistent Force Field Captures Homologue-Resolved HP1 Phase Separation. Journal of Chemical Theory and Computation 2021, 17, 3134–3144.

(35) Tesei, G.; Schulze, T. K.; Crehuet, R.; Lindorff-Larsen, K. Accurate Model of Liquid–Liquid Phase Behavior of Intrinsically Disordered Proteins from Optimization of Single-Chain Properties. Proceedings of the National Academy of Sciences 2021, 118, e2111696118.

(36) Benayad, Z.; von Bülow, S.; Stelzl, L. S.; Hummer, G. Simulation of FUS Protein Condensates with an Adapted Coarse-Grained Model. Journal of Chemical Theory and Computation 2021, 17, 525–537.

(37) Farag, M.; Borcherds, W. M.; Bremer, A.; Mittag, T.; Pappu, R. V. Phase Separation of Protein Mixtures Is Driven by the Interplay of Homotypic and Heterotypic Interactions. Nature Communications 2023, 14, 5527.

(38) Lotthammer, J. M.; Ginell, G. M.; Griffith, D.; Emenecker, R. J.; Holehouse, A. S. Direct Prediction of Intrinsically Disordered Protein Conformational Properties from Sequence. Nature Methods 2024,

(39) von Bülow, S.; Tesei, G.; Lindorff-Larsen, K. Prediction of Phase Separation Propensities of Disordered Proteins from Sequence. 2024.

(40) Zhang, Y.; Li, S.; Gong, X.; Chen, J. Toward Accurate Simulation of Coupling between Protein Secondary Structure and Phase Separation. Journal of the American Chemical Society 2024, 146, 342–357.

(41) Kapoor, U.; Kim, Y. C.; Mittal, J. Coarse-Grained Models to Study Protein–DNA Interactions and Liquid–Liquid Phase Separation. Journal of Chemical Theory and Computation 2024, 20, 1717–1731.

(42) Liu, S.; Wang, C.; Latham, A. P.; Ding, X.; Zhang, B. OpenABC Enables Flexible, Simplified, and Efficient GPU Accelerated Simulations of Biomolecular Condensates. PLOS Computational Biology 2023, 19, e1011442.

(43) Wang, J.; Choi, J.-M.; Holehouse, A. S.; Lee, H. O.; Zhang, X.; Jahnel, M.; Maharana, S.; Lemaitre, R.; Pozniakovsky, A.; Drechsel, D.; Poser, I.; Pappu, R. V.; Alberti, S.; Hyman, A. A. A Molecular Grammar Governing the Driving Forces for Phase Separation of Prion-like RNA Binding Proteins. Cell 2018, 174, 688–699.e16.

(44) Prusty, D.; Pryamitsyn, V.; Olvera de la Cruz, M. Thermodynamics of Associative Polymer Blends. Macromolecules 2018, 51, 5918–5932.

(45) Choi, J.-M.; Dar, F.; Pappu, R. V. LASSI: A Lattice Model for Simulating Phase Transitions of Multivalent Proteins. PLOS Computational Biology 2019, 15, e1007028.

(46) Zhang, Y.; Xu, B.; Weiner, B. G.; Meir, Y.; Wingreen, N. S. Decoding the Physical Principles of Two-Component Biomolecular Phase Separation. eLife 2021, 10, e62403.

(47) Harmon, T. S.; Holehouse, A. S.; Rosen, M. K.; Pappu, R. V. Intrinsically Disordered Linkers Determine the Interplay between Phase Separation and Gelation in Multivalent Proteins. eLife 6, e30294.

(48) Bremer, A.; Farag, M.; Borcherds, W. M.; Peran, I.; Martin, E. W.; Pappu, R. V.; Mittag, T. Deciphering How Naturally Occurring Sequence Features Impact the Phase Behaviours of Disordered Prion-like Domains. Nature Chemistry 2022, 14, 196–207.

(49) Farag, M.; Cohen, S. R.; Borcherds, W. M.; Bremer, A.; Mittag, T.; Pappu, R. V. Condensates Formed by Prion-like Low-Complexity Domains Have Small-World Network Structures and Interfaces Defined by Expanded Conformations. Nature Communications 2022, 13, 7722.

(50) Ranganathan, S.; Shakhnovich, E. I. Dynamic Metastable Long-Living Droplets Formed by Sticker-Spacer Proteins. eLife 2020, 9, e56159.

(51) Sood, A.; Zhang, B. Preserving Condensate Structure and Composition by Lowering Sequence Complexity. Biophysical Journal 2024, 123, 1815–1826.

(52) Riley, A. C.; Ashlock, D. A.; Graether, S. P. The Difficulty of Aligning Intrinsically Disordered Protein Sequences as Assessed by Conservation and Phylogeny. PLOS ONE 2023, 18, e0288388.

(53) Oldfield, C. J.; Dunker, A. K. Intrinsically Disordered Proteins and Intrinsically Disordered Protein Regions. Annual Review of Biochemistry 2014, 83, 553–584.

(54) Brown, C. J.; Takayama, S.; Campen, A. M.; Vise, P.; Marshall, T. W.; Oldfield, C. J.; Williams, C. J.; Dunker, A. K. Evolutionary Rate Heterogeneity in Proteins with Long Disordered Regions. Journal of Molecular Evolution 2002, 55, 104–110.

(55) Huang, H.; Sarai, A. Analysis of the Relationships between Evolvability, Thermodynamics, and the Functions of Intrinsically Disordered Proteins/Regions. Computational Biology and Chemistry 2012, 41, 51–57.

(56) Nunez-Castilla, J.; Siltberg-Liberles, J. In Intrinsically Disordered Proteins: Methods and Protocols; Kragelund, B. B., Skriver, K., Eds.; Springer US: New York, NY, 2020; pp 147–177.

(57) Thompson, J. D.; Linard, B.; Lecompte, O.; Poch, O. A Comprehensive Benchmark Study of Multiple Sequence Alignment Methods: Current Challenges and Future Perspectives. PLOS ONE 2011, 6, e18093.

(58) Lange, J.; Wyrwicz, L. S.; Vriend, G. KMAD: Knowledge-Based Multiple Sequence Alignment for Intrinsically Disordered Proteins. Bioinformatics 2016, 32, 932–936.

(59) Dasmeh, P.; Doronin, R.; Wagner, A. The Length Scale of Multivalent Interactions Is Evolutionarily Conserved in Fungal and Vertebrate Phase-Separating Proteins. Genetics 2022, 220, iyab184.

(60) Lu, A. X.; Lu, A. X.; Pritišanac, I.; Zarin, T.; Forman-Kay, J. D.; Moses, A. M. Discovering Molecular Features of Intrinsically Disordered Regions by Using Evolution for Contrastive Learning. PLOS Computational Biology 2022, 18, e1010238.

(61) Ho, W.-L.; Huang, J.-r. The Return of the Rings: Evolutionary Convergence of Aromatic Residues in the Intrinsically Disordered Regions of RNA-binding Proteins for Liquid–Liquid Phase Separation. Protein Science 2022, 31, e4317.

(62) Ofer, D.; Brandes, N.; Linial, M. The Language of Proteins: NLP, Machine Learning & Protein Sequences. Computational and Structural Biotechnology Journal 2021, 19, 1750–1758.

(63) Rives, A.; Meier, J.; Sercu, T.; Goyal, S.; Lin, Z.; Liu, J.; Guo, D.; Ott, M.; Zitnick, C. L.; Ma, J.; Fergus, R. Biological Structure and Function Emerge from Scaling Unsupervised Learning to 250 Million Protein Sequences. Proceedings of the National Academy of Sciences 2021, 118, e2016239118.

(64) Elnaggar, A.; Ding, W.; Jones, L.; Gibbs, T.; Feher, T.; Angerer, C.; Severini, S.; Matthes, F.; Rost, B. CodeTrans: Towards Cracking the Language of Silicon’s Code Through Self-Supervised Deep Learning and High Performance Computing. 2021.

(65) Strodthoff, N.; Wagner, P.; Wenzel, M.; Samek, W. UDSMProt: Universal Deep Sequence Models for Protein Classification. Bioinformatics 2020, 36, 2401–2409.

(66) Alley, E. C.; Khimulya, G.; Biswas, S.; AlQuraishi, M.; Church, G. M. Unified Rational Protein Engineering with Sequence-Based Deep Representation Learning. Nature Methods 2019, 16, 1315–1322.

(67) Brandes, N.; Ofer, D.; Peleg, Y.; Rappoport, N.; Linial, M. ProteinBERT: A Universal Deep-Learning Model of Protein Sequence and Function. Bioinformatics 2022, 38, 2102–2110.

(68) Boutet, E.; Lieberherr, D.; Tognolli, M.; Schneider, M.; Bansal, P.; Bridge, A. J.; Poux, S.; Bougueleret, L.; Xenarios, I. In Plant Bioinformatics: Methods and Protocols; Edwards, D., Ed.; Springer: New York, NY, 2016; pp 23–54.

(69) Vaswani, A.; Shazeer, N.; Parmar, N.; Uszkoreit, J.; Jones, L.; Gomez, A. N.; Kaiser, Ł.; Polosukhin, I. Attention Is All You Need. Advances in Neural Information Processing Systems. 2017.

(70) Meier, J.; Rao, R.; Verkuil, R.; Liu, J.; Sercu, T.; Rives, A. Language Models Enable Zero-Shot Prediction of the Effects of Mutations on Protein Function. Proceedings of the 35th International Conference on Neural Information Processing Systems. Red Hook, NY, USA, 2024; pp 29287–29303.

(71) Cagiada, M.; Jonsson, N.; Lindorff-Larsen, K. Decoding Molecular Mechanisms for Loss of Function Variants in the Human Proteome. 2024.

(72) Ouyang-Zhang, J.; Klivans, A. R.; Diaz, D. J.; Krähenbühl, P. Predicting a Protein’s Stability under a Million Mutations.

(73) Lin, W.; Wells, J.; Wang, Z.; Orengo, C.; Martin, A. C. R. Enhancing Missense Variant Pathogenicity Prediction with Protein Language Models Using VariPred. Scientific Reports 2024, 14, 8136.

(74) Saadat, A.; Fellay, J. Fine-Tuning the ESM2 Protein Language Model to Understand the Functional Impact of Missense Variants. Computational and Structural Biotechnology Journal 2025, 27, 2199–2207.

(75) Chu, S. K. S.; Narang, K.; Siegel, J. B. Protein Stability Prediction by Fine-Tuning a Protein Language Model on a Mega-Scale Dataset. PLOS Computational Biology 2024, 20, e1012248.

(76) Marquet, C.; Schlensok, J.; Abakarova, M.; Rost, B.; Laine, E. Expert-Guided Protein Language Models Enable Accurate and Blazingly Fast Fitness Prediction. Bioinformatics 2024, 40, btae621.

(77) Gong, J.; Jiang, L.; Chen, Y.; Zhang, Y.; Li, X.; Ma, Z.; Fu, Z.; He, F.; Sun, P.; Ren, Z.; Tian, M. THPLM: A Sequence-Based Deep Learning Framework for Protein Stability Changes Prediction upon Point Variations Using Pretrained Protein Language Model. Bioinformatics 2023, 39, btad646.

(78) Brandes, N.; Goldman, G.; Wang, C. H.; Ye, C. J.; Ntranos, V. Genome-Wide Prediction of Disease Variant Effects with a Deep Protein Language Model. Nature Genetics 2023, 55, 1512–1522.

(79) Lin, Z.; Akin, H.; Rao, R.; Hie, B.; Zhu, Z.; Lu, W.; Smetanin, N.; Verkuil, R.; Kabeli, O.; Shmueli, Y.; Fazel-Zarandi, M.; Sercu, T.; Candido, S.; Rives, A. Evolutionary-Scale Prediction of Atomic-Level Protein Structure with a Language Model. 2023,

(80) Zeng, W.; Dou, Y.; Pan, L.; Xu, L.; Peng, S. Improving Prediction Performance of General Protein Language Model by Domain-Adaptive Pretraining on DNA-binding Protein. Nature Communications 2024, 15, 7838.

(81) Highly Accurate Protein Structure Prediction with AlphaFold | Nature. https://www.nature.com/articles/s41586-021-03819-2#citeas.

(82) Ruff, K. M.; Pappu, R. V. AlphaFold and Implications for Intrinsically Disordered Proteins. Journal of Molecular Biology 2021, 433, 167208.

(83) Alderson, T. R.; Pritišanac, I.; Kolarić, Đ.; Moses, A. M.; Forman-Kay, J. D. Systematic Identification of Conditionally Folded Intrinsically Disordered Regions by AlphaFold2. Proceedings of the National Academy of Sciences of the United States of America 120, e2304302120.

(84) Larson, A. G.; Elnatan, D.; Keenen, M. M.; Trnka, M. J.; Johnston, J. B.; Burlingame, A. L.; Agard, D. A.; Redding, S.; Narlikar, G. J. Liquid Droplet Formation by HP1α Suggests a Role for Phase Separation in Heterochromatin. Nature 2017, dirusso2023, 236–240.

(85) Sanulli, S.; Trnka, M. J.; Dharmarajan, V.; Tibble, R. W.; Pascal, B. D.; Burlingame, A. L.; Griffin, P. R.; Gross, J. D.; Narlikar, G. J. HP1 Reshapes Nucleosome Core to Promote Phase Separation of Heterochromatin. Nature 2019, 575, 390–394.

(86) Tortora, M. M. C.; Brennan, L. D.; Karpen, G.; Jost, D. HP1-driven Phase Separation Recapitulates the Thermodynamics and Kinetics of Heterochromatin Condensate Formation. Proceedings of the National Academy of Sciences 2023, 120, e2211855120.

(87) Brennan, L.; Kim, H.-K.; Colmenares, S.; Ego, T.; Kumar, A.; Safran, S.; Ryu, J.-K.; Karpen, G. HP1a Promotes Chromatin Liquidity and Drives Spontaneous Heterochromatin Compartmentalization. 2024.

(88) Smothers, J. F.; Henikoff, S. The Hinge and Chromo Shadow Domain Impart Distinct Targeting of HP1-like Proteins. Molecular and Cellular Biology 2001, 21, 2555–2569.

(89) Badugu, R.; Yoo, Y.; Singh, P. B.; Kellum, R. Mutations in the Heterochromatin Protein 1 (HP1) Hinge Domain Affect HP1 Protein Interactions and Chromosomal Distribution. Chromosoma 2005, 113, 370–384.

(90) Strom, A. R. et al. HP1α Is a Chromatin Crosslinker That Controls Nuclear and Mitotic Chromosome Mechanics. eLife 2021, 10, e63972.

(91) Shakhnovich, E.; Abkevich, V.; Ptitsyn, O. Conserved Residues and the Mechanism of Protein Folding. Nature 1996, 379, 96–98.

(92) Hamill, S. J.; Cota, E.; Chothia, C.; Clarke, J. Conservation of Folding and Stability within a Protein Family: The Tyrosine Corner as an Evolutionary Cul-de-Sac1. Journal of Molecular Biology 2000, 295, 641–649.

(93) Ingles-Prieto, A.; Ibarra-Molero, B.; Delgado-Delgado, A.; Perez-Jimenez, R.; Fernandez, J. M.; Gaucher, E. A.; Sanchez-Ruiz, J. M.; Gavira, J. A. Conservation of Protein Structure over Four Billion Years. Structure (London, England : 1993) 2013, 21, 1690–1697.

(94) Brown, C. J.; Johnson, A. K.; Dunker, A. K.; Daughdrill, G. W. Evolution and Disorder. Current opinion in structural biology 2011, 21, 441–446.

(95) Liu, Z.; Huang, Y. Advantages of Proteins Being Disordered. Protein Science 2014, 23, 539–550.

(96) Forman-Kay, J. D.; Mittag, T. From Sequence and Forces to Structure, Function, and Evolution of Intrinsically Disordered Proteins. Structure 2013, 21, 1492–1499.

(97) Sharma, R.; Kumar, S.; Tsunoda, T.; Patil, A.; Sharma, A. Predicting MoRFs in Protein Sequences Using HMM Profiles. BMC Bioinformatics 2016, 17, 504.

(98) Jarnot, P.; Ziemska-Legiecka, J.; Grynberg, M.; Gruca, A. Insights from Analyses of Low Complexity Regions with Canonical Methods for Protein Sequence Comparison. Briefings in Bioinformatics 2022, 23, bbac299.

(99) Peng, Z.; Li, Z.; Meng, Q.; Zhao, B.; Kurgan, L. CLIP: Accurate Prediction of Disordered Linear Interacting Peptides from Protein Sequences Using Co-Evolutionary Information. Briefings in Bioinformatics 2023, 24, bbac502.

(100) Sagawa, T.; Kanao, E.; Ogata, K.; Imami, K.; Ishihama, Y. Prediction of Protein Half-lives from Amino Acid Sequences by Protein Language Models. 2024.

(101) Yang, J.; Cheng, W.-x.; Wu, G.; Sheng, S.; Zhang, P. Prediction of Folding Patterns for Intrinsic Disordered Protein. Scientific Reports 2023, 13, 20343.

(102) Chavali, S.; Singh, A. K.; Santhanam, B.; Babu, M. M. Amino Acid Homorepeats in Proteins. Nature Reviews. Chemistry 2020, 4, 420–434.

(103) Farahi, N.; Lazar, T.; Wodak, S. J.; Tompa, P.; Pancsa, R. Integration of Data from Liquid–Liquid Phase Separation Databases Highlights Concentration and Dosage Sensitivity of LLPS Drivers. International Journal of Molecular Sciences 2021, 22, 3017.

(104) Orti, F.; Fernández, M. L.; Marino-Buslje, C. MLOsMetaDB, a Meta-Database to Centralize the Information on Liquid–Liquid Phase Separation Proteins and Membraneless Organelles. Protein Science 2024, 33, e4858.

(105) Mészáros, B.; Erdős, G.; Szabó, B.; Schád, É.; Tantos, Á.; Abukhairan, R.; Horváth, T.; Murvai, N.; Kovács, O. P.; Kovács, M.; Tosatto, S. C. E.; Tompa, P.; Dosztányi, Z.; Pancsa, R. PhaSePro: The Database of Proteins Driving Liquid–Liquid Phase Separation. Nucleic Acids Research 2020, 48, D360–D367.

(106) You, K.; Huang, Q.; Yu, C.; Shen, B.; Sevilla, C.; Shi, M.; Hermjakob, H.; Chen, Y.; Li, T. PhaSepDB: A Database of Liquid–Liquid Phase Separation Related Proteins. Nucleic Acids Research 2020, 48, D354–D359.

(107) Ning, W.; Guo, Y.; Lin, S.; Mei, B.; Wu, Y.; Jiang, P.; Tan, X.; Zhang, W.; Chen, G.; Peng, D.; Chu, L.; Xue, Y. DrLLPS: A Data Resource of Liquid–Liquid Phase Separation in Eukaryotes. Nucleic Acids Research 2020, 48, D288–D295.

(108) Li, Q.; Peng, X.; Li, Y.; Tang, W.; Zhu, J.; Huang, J.; Qi, Y.; Zhang, Z. LLPSDB: A Database of Proteins Undergoing Liquid–Liquid Phase Separation in Vitro. Nucleic Acids Research 2020, 48, D320–D327.

(109) Saar, K. L.; Morgunov, A. S.; Qi, R.; Arter, W. E.; Krainer, G.; Lee, A. A.; Knowles, T. P. J. Learning the Molecular Grammar of Protein Condensates from Sequence Determinants and Embeddings. Proceedings of the National Academy of Sciences 2021, 118, e2019053118.

(110) Ozawa, Y.; Anbo, H.; Ota, M.; Fukuchi, S. Classification of Proteins Inducing Liquid–Liquid Phase Separation: Sequential, Structural and Functional Characterization. Journal of Biochemistry 2022, 173, 255–264.

(111) Rekhi, S.; Garcia, C. G.; Barai, M.; Rizuan, A.; Schuster, B. S.; Kiick, K. L.; Mittal, J. Expanding the Molecular Language of Protein Liquid–Liquid Phase Separation. Nature Chemistry 2024, 16, 1113–1124.

(112) Li, S.; Zhang, Y.; Chen, J. Backbone Interactions and Secondary Structures in Phase Separation of Disordered Proteins. Biochemical Society Transactions 2024, 52, 319–329.

(113) Schuster, B. S.; Dignon, G. L.; Tang, W. S.; Kelley, F. M.; Ranganath, A. K.; Jahnke, C. N.; Simpkins, A. G.; Regy, R. M.; Hammer, D. A.; Good, M. C.; Mittal, J. Identifying Sequence Perturbations to an Intrinsically Disordered Protein That Determine Its Phase-Separation Behavior. Proceedings of the National Academy of Sciences of the United States of America 2020, 117, 11421–11431.

(114) Pak, C. W.; Kosno, M.; Holehouse, A. S.; Padrick, S. B.; Mittal, A.; Ali, R.; Yunus, A. A.; Liu, D. R.; Pappu, R. V.; Rosen, M. K. Sequence Determinants of Intracellular Phase Separation by Complex Coacervation of a Disordered Protein. Molecular Cell 2016, 63, 72–85.

(115) Cohan, M. C.; Shinn, M. K.; Lalmansingh, J. M.; Pappu, R. V. Uncovering Nonrandom Binary Patterns Within Sequences of Intrinsically Disordered Proteins. Journal of Molecular Biology 2022, 434, 167373.

(116) Dao, T. P.; Kolaitis, R.-M.; Kim, H. J.; O’Donovan, K.; Martyniak, B.; Colicino, E.; Hehnly, H.; Taylor, J. P.; Castañeda, C. A. Ubiquitin Modulates Liquid-Liquid Phase Separation of UBQLN2 via Disruption of Multivalent Interactions. Molecular cell 2018, 69, 965–978.e6.

(117) Tatavosian, R.; Kent, S.; Brown, K.; Yao, T.; Duc, H. N.; Huynh, T. N.; Zhen, C. Y.; Ma, B.; Wang, H.; Ren, X. Nuclear Condensates of the Polycomb Protein Chromobox 2 (CBX2) Assemble through Phase Separation. Journal of Biological Chemistry 2019, 294, 1451–1463.

(118) Yang, Y.; Jones, H. B.; Dao, T. P.; Castañeda, C. A. Single Amino Acid Substitutions in Stickers, but Not Spacers, Substantially Alter UBQLN2 Phase Transitions and Dense Phase Material Properties. The Journal of Physical Chemistry B 2019, 123, 3618–3629.

(119) DiRusso, C. J.; DeMaria, A. M.; Wong, J.; Wang, W.; Jordanides, J. J.; Whitty, A.; Allen, K. N.; Gilmore, T. D. A Conserved Core Region of the Scaffold NEMO Is Essential for Signal-Induced Conformational Change and Liquid-Liquid Phase Separation. Journal of Biological Chemistry 2023, 299, 105396.

(120) Mitrea, D. M.; Cika, J. A.; Guy, C. S.; Ban, D.; Banerjee, P. R.; Stanley, C. B.; Nourse, A.; Deniz, A. A.; Kriwacki, R. W. Nucleophosmin Integrates within the Nucleolus via Multi-Modal Interactions with Proteins Displaying R-rich Linear Motifs and rRNA. eLife 2016, 5, e13571.

(121) Xiao, Y.; Chen, J.; Wan, Y.; Gao, Q.; Jing, N.; Zheng, Y.; Zhu, X. Regulation of Zebrafish Dorsoventral Patterning by Phase Separation of RNA-binding Protein Rbm14. Cell Discovery 2019, 5, 1–17.

(122) Mitrea, D. M.; Cika, J. A.; Stanley, C. B.; Nourse, A.; Onuchic, P. L.; Banerjee, P. R.; Phillips, A. H.; Park, C.-G.; Deniz, A. A.; Kriwacki, R. W. Self-Interaction of NPM1 Modulates Multiple Mechanisms of Liquid–Liquid Phase Separation. Nature Communications 2018, 9, 842.

(123) Wang, B.; Zhang, L.; Dai, T.; Qin, Z.; Lu, H.; Zhang, L.; Zhou, F. Liquid–Liquid Phase Separation in Human Health and Diseases. Signal Transduction and Targeted Therapy 2021, 6, 1–16.

(124) Landrum, M. J.; Lee, J. M.; Riley, G. R.; Jang, W.; Rubinstein, W. S.; Church, D. M.; Maglott, D. R. ClinVar: Public Archive of Relationships among Sequence Variation and Human Phenotype. Nucleic Acids Research 2014, 42, D980–D985.

(125) Landrum, M. J. et al. ClinVar: Improving Access to Variant Interpretations and Supporting Evidence. Nucleic Acids Research 2018, 46, D1062–D1067.

(126) Coudert, E.; Gehant, S.; de Castro, E.; Pozzato, M.; Baratin, D.; Neto, T.; Sigrist, C. J. A.; Redaschi, N.; Bridge, A.; The UniProt Consortium Annotation of Biologically Relevant Ligands in UniProtKB Using ChEBI. Bioinformatics 2023, 39, btac793.

(127) Rostam, N.; Ghosh, S.; Chow, C. F. W.; Hadarovich, A.; Landerer, C.; Ghosh, R.; Moon, H.; Hersemann, L.; Mitrea, D. M.; Klein, I. A.; Hyman, A. A.; Toth-Petroczy, A. CD-CODE: Crowdsourcing Condensate Database and Encyclopedia. Nature Methods 2023, 20, 673–676.

(128) Varadi, M. et al. AlphaFold Protein Structure Database in 2024: Providing Structure Coverage for over 214 Million Protein Sequences. Nucleic Acids Research 2024, 52, D368–D375.

(129) Suzek, B. E.; Huang, H.; McGarvey, P.; Mazumder, R.; Wu, C. H. UniRef: Comprehensive and Non-Redundant UniProt Reference Clusters. Bioinformatics 2007, 23, 1282–1288.

(130) Suzek, B. E.; Wang, Y.; Huang, H.; McGarvey, P. B.; Wu, C. H. UniRef Clusters: A Comprehensive and Scalable Alternative for Improving Sequence Similarity Searches. Bioinformatics 2015, 31, 926–932.

(131) Remmert, M.; Biegert, A.; Hauser, A.; Söding, J. HHblits: Lightning-Fast Iterative Protein Sequence Searching by HMM-HMM Alignment. Nature Methods 2012, 9, 173–175.

(132) Steinegger, M.; Meier, M.; Mirdita, M.; Vöhringer, H.; Haunsberger, S. J.; Söding, J. HH-suite3 for Fast Remote Homology Detection and Deep Protein Annotation. BMC Bioinformatics 2019, 20, 473.

(133) Mirdita, M.; von den Driesch, L.; Galiez, C.; Martin, M. J.; Söding, J.; Steinegger, M. Uniclust Databases of Clustered and Deeply Annotated Protein Sequences and Alignments. Nucleic Acids Research 2017, 45, D170–D176.

