## Supplementary Information for "Protein Language Model Identifies Disordered, Conserved Motifs Implicated in Phase Separation"

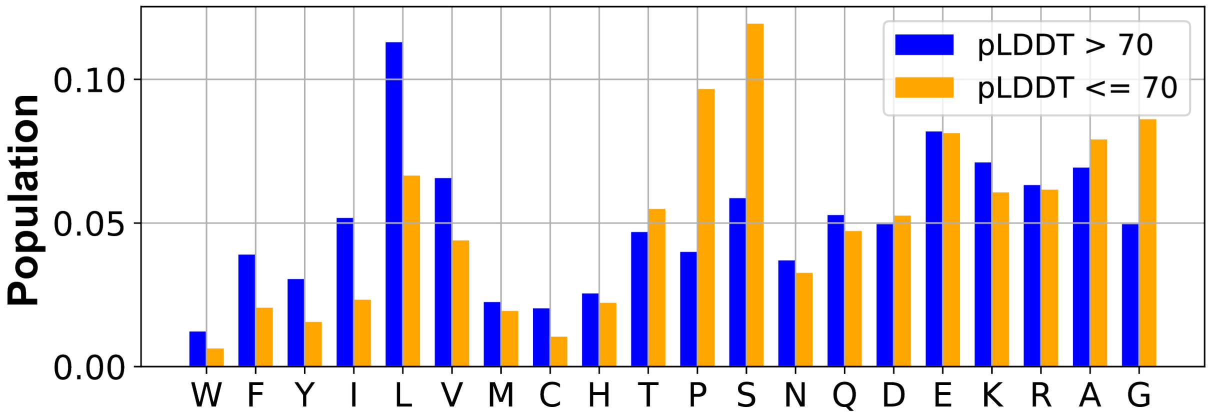


**Figure S1: Fraction of different amino acids in structured and disordered residues identified from proteins in the MLO-hProt dataset (939).** The structured (pLDDT >70) and the disordered (pLDDT <= 70) residues were identified using the AlphaFold2 pLDDT score.

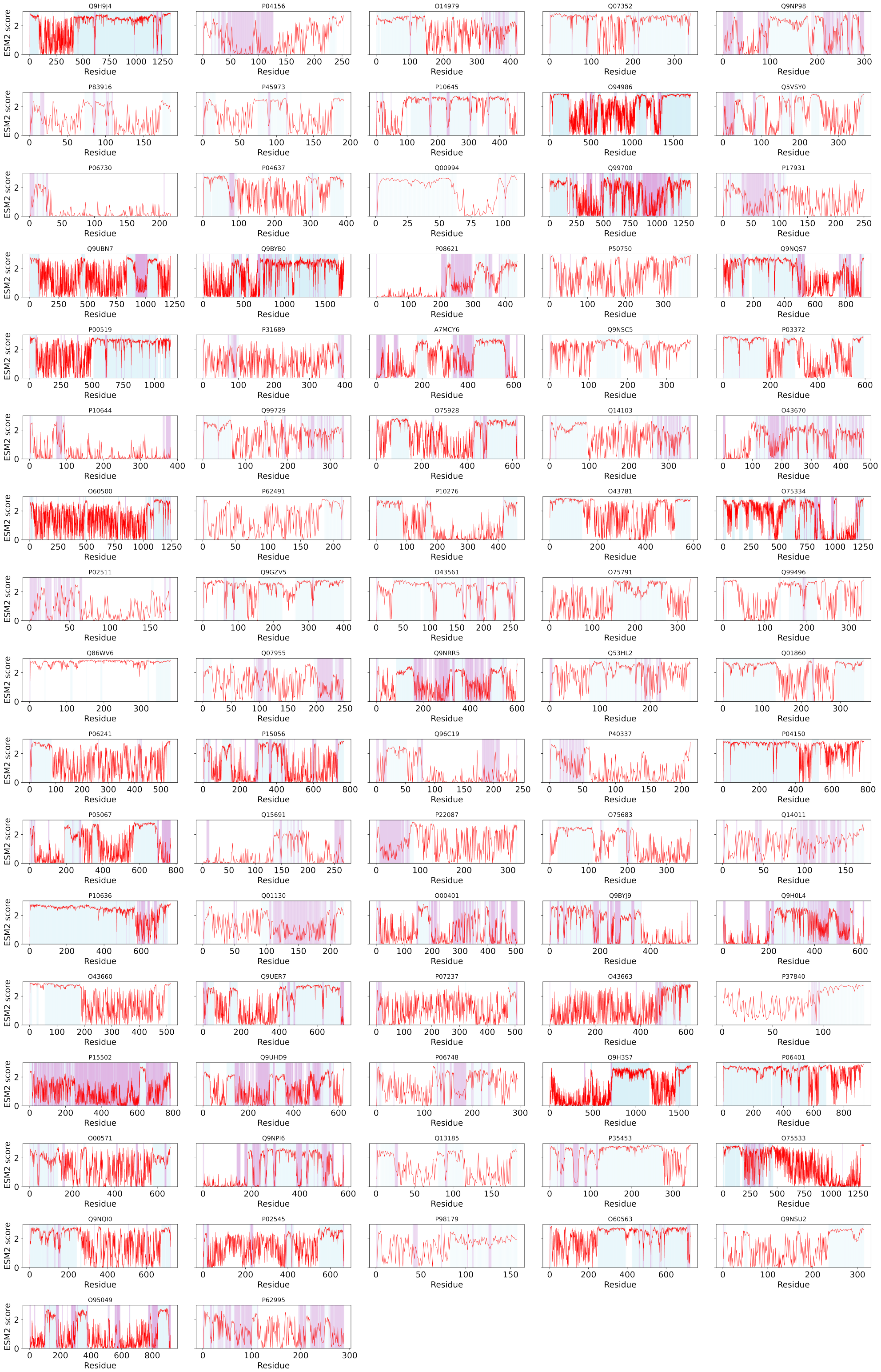


**Figure S2: ESM2 and AlphaFold2 predictions for all proteins in the dMLO-hProt dataset.** ESM2 scores for all amino acids in the dMLO-hIDR proteins (full-length) are shown, with UniProt IDs indicated above each plot. Residues with pLDDT scores $\leq 70$ are highlighted in blue, indicating regions lacking well-defined structure. Of these residues, those with ESM2 scores $\leq 1.5$ are colored plum, marking conserved disordered segments.

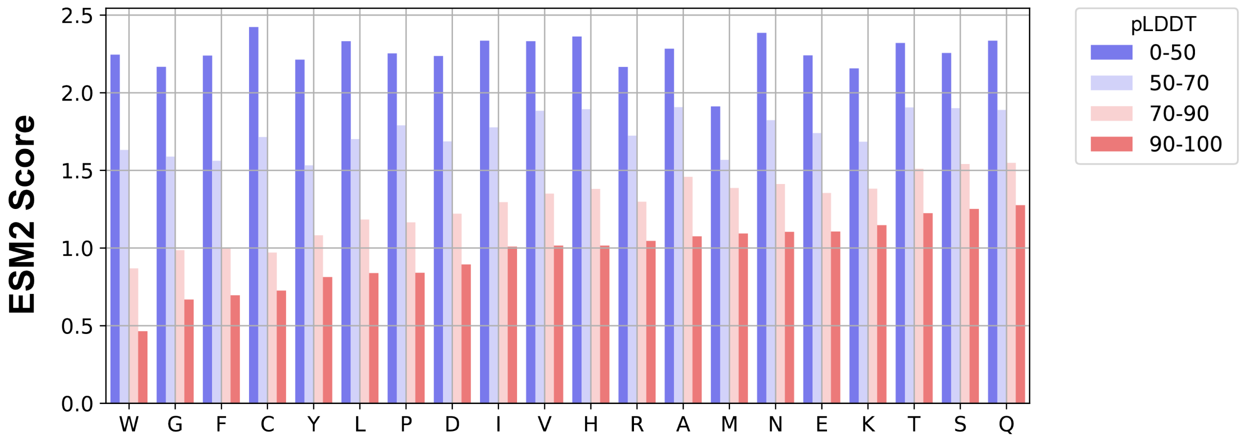


**Figure S3: Average ESM2 score for amino acids with different structural order, as indicated by the colors.** The averaging was performed over residues from proteins in the MLO-hProt dataset (939).

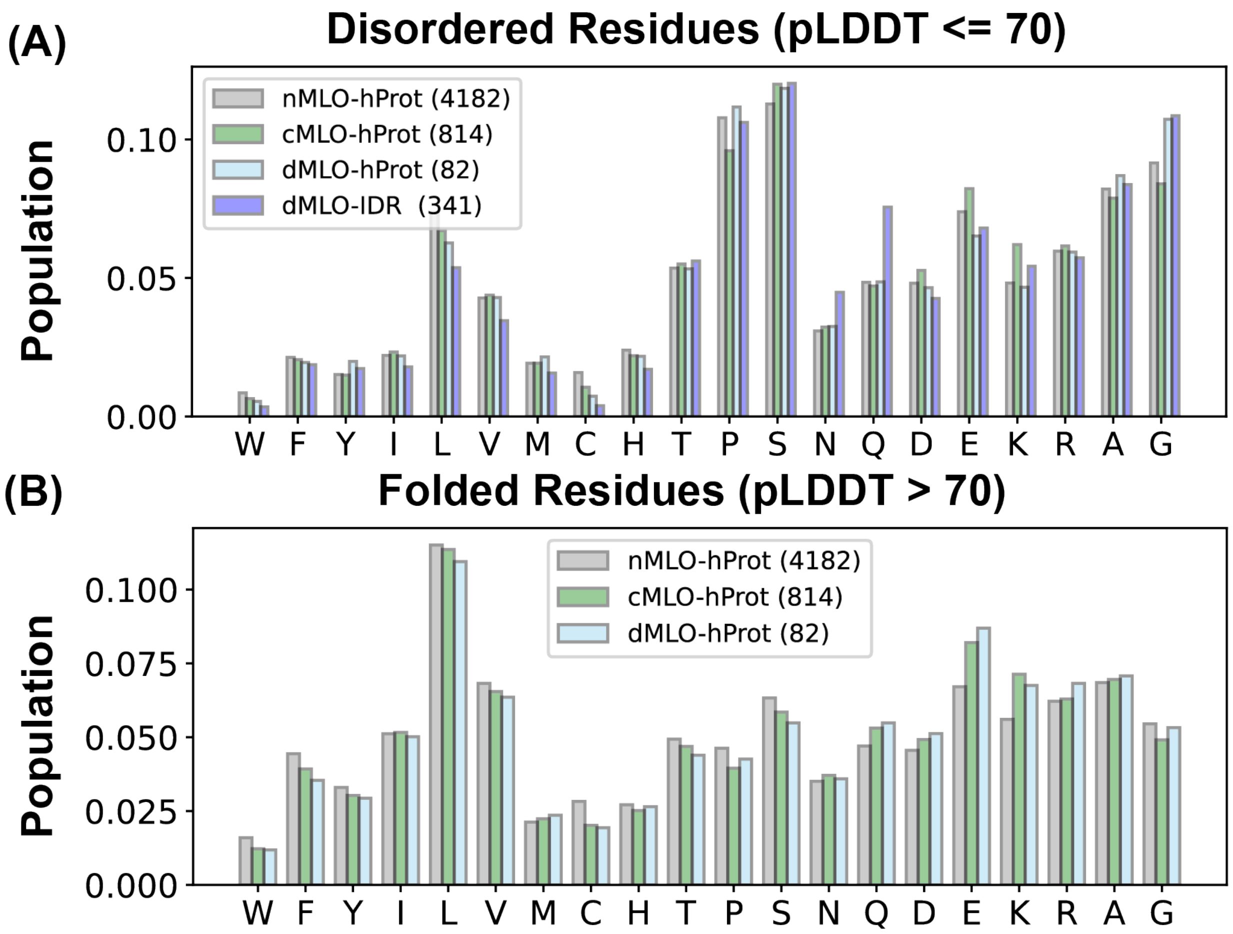


**Figure S4: Fraction of different amino acids across proteins from various datasets.** Panels (A) and (B) depict the amino acid composition of disordered and folded residues, respectively. For proteins in the nMLO-hProt, cMLO-hProt, and dMLO-hProt datasets, residue structural order was assessed using the AlphaFold2 pLDDT score. In contrast, disordered residues for proteins in the dMLO-IDR dataset were obtained from the MLOsmetaDB database.[S1]

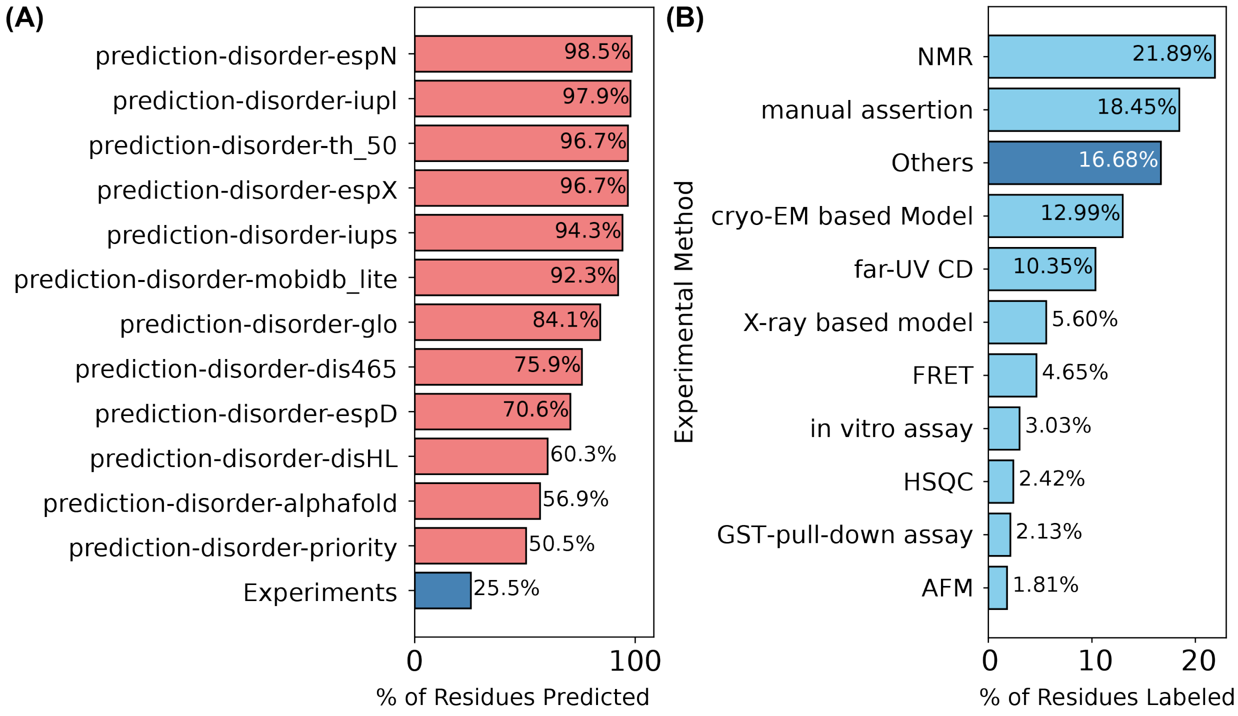


**Figure S5: Disorder annotation of dMLO-IDR proteins in the MobiDB database.**[S2] (A) Computational tools used for predicting amino acid disorder. The numerical values represent the fraction of amino acids annotated as disordered by each tool. (B) Experimental techniques used to determine amino acid disorder. The numerical values represent the fraction of amino acids identified as disordered by each method.

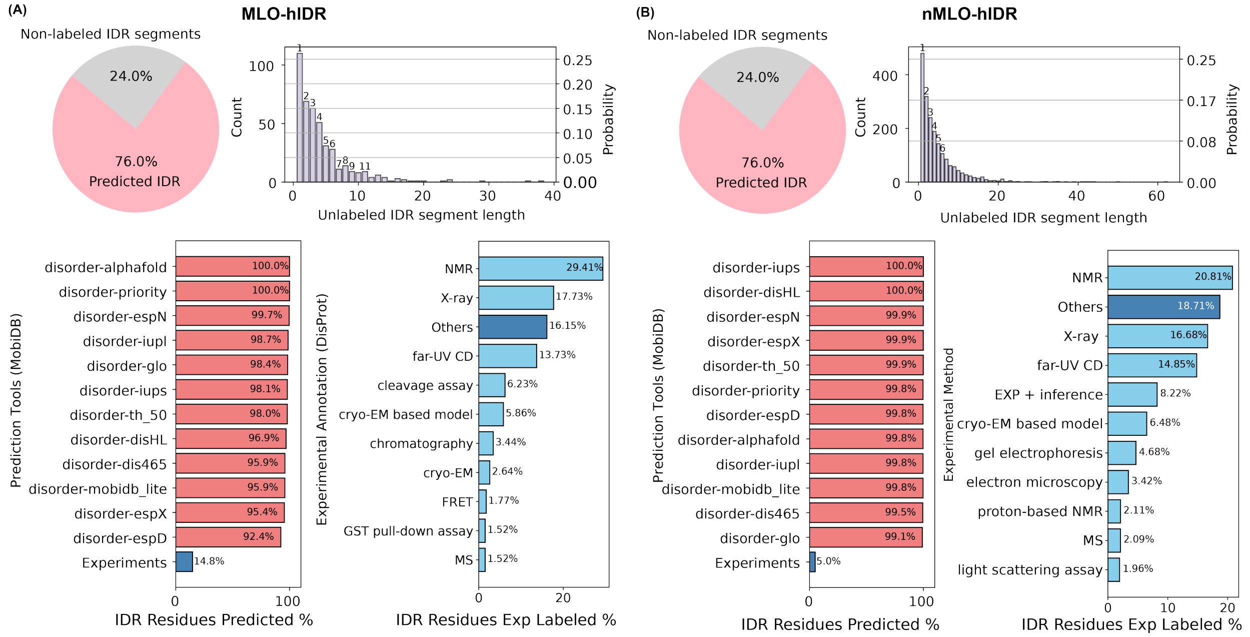


**Figure S6: Disorder annotation of human proteins in the MobiDB database.**[S2] (A) The top left panel shows a pie chart depicting the fraction of amino acids annotated as disordered (pink) and not annotated (grey) by the MobiDB database within the MLO-hIDR dataset. The top right panel presents a histogram of the lengths of the 24% of MLO-hIDRs lacking disorder annotation. The bottom two plots display distributions analogous to those in Figure S5. (B) Same as in panel A, but for the nMLO-hIDR dataset.

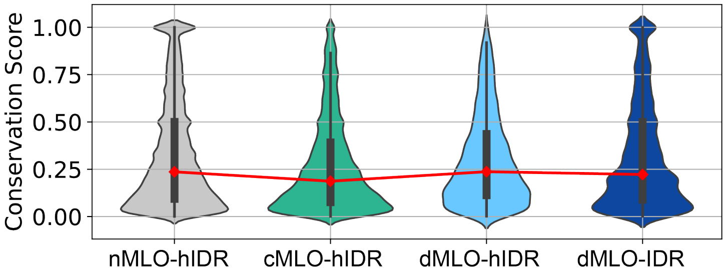


**Figure S7: Violin plots illustrating the distribution of conservation scores for disordered residues across the nMLO–hIDR, cMLO–hIDR, dMLO–hIDR, and dMLO–IDR datasets.** Pairwise statistical comparisons were conducted using two-sided Mann–Whitney U tests on the conservation score distributions (null hypothesis: the two groups have equal medians). P-values indicate the probability of observing the observed rank differences under the null hypothesis. Statistical significance is denoted as follows: ***: $p < 0.001$; **: $p < 0.01$; *:$p < 0.05$;

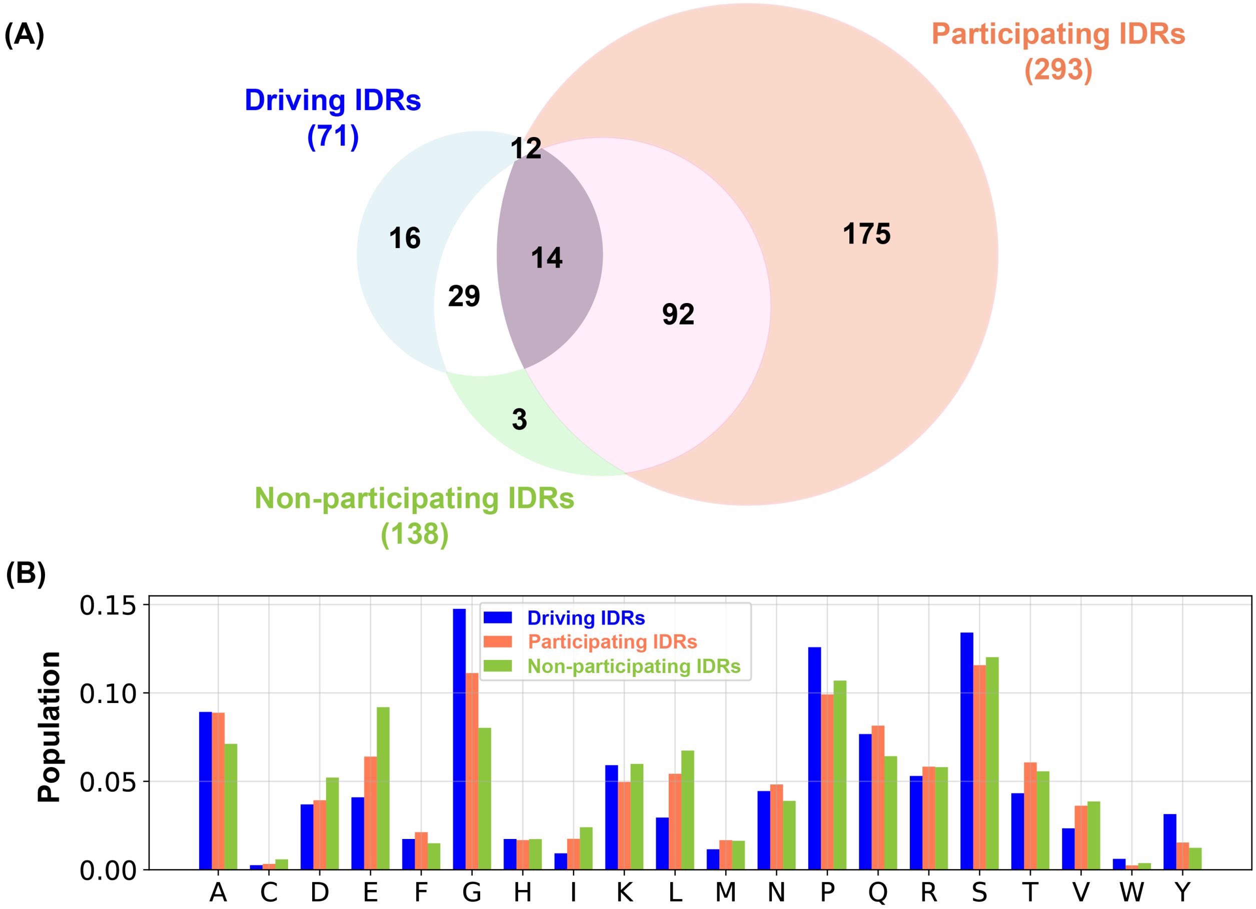


**Figure S8: Statistics of the three subgroups of IDRs in the dMLO-IDR dataset.** (A) The numbers below the labels, as well as the areas of the circles with corresponding colors, represent the total count of IDR segments in each subgroup. Since IDR segments from distinct subgroups may originate from the same protein, overlaps between the circles are observed. (B) Proportion of amino acids across IDRs in the three subgroups.

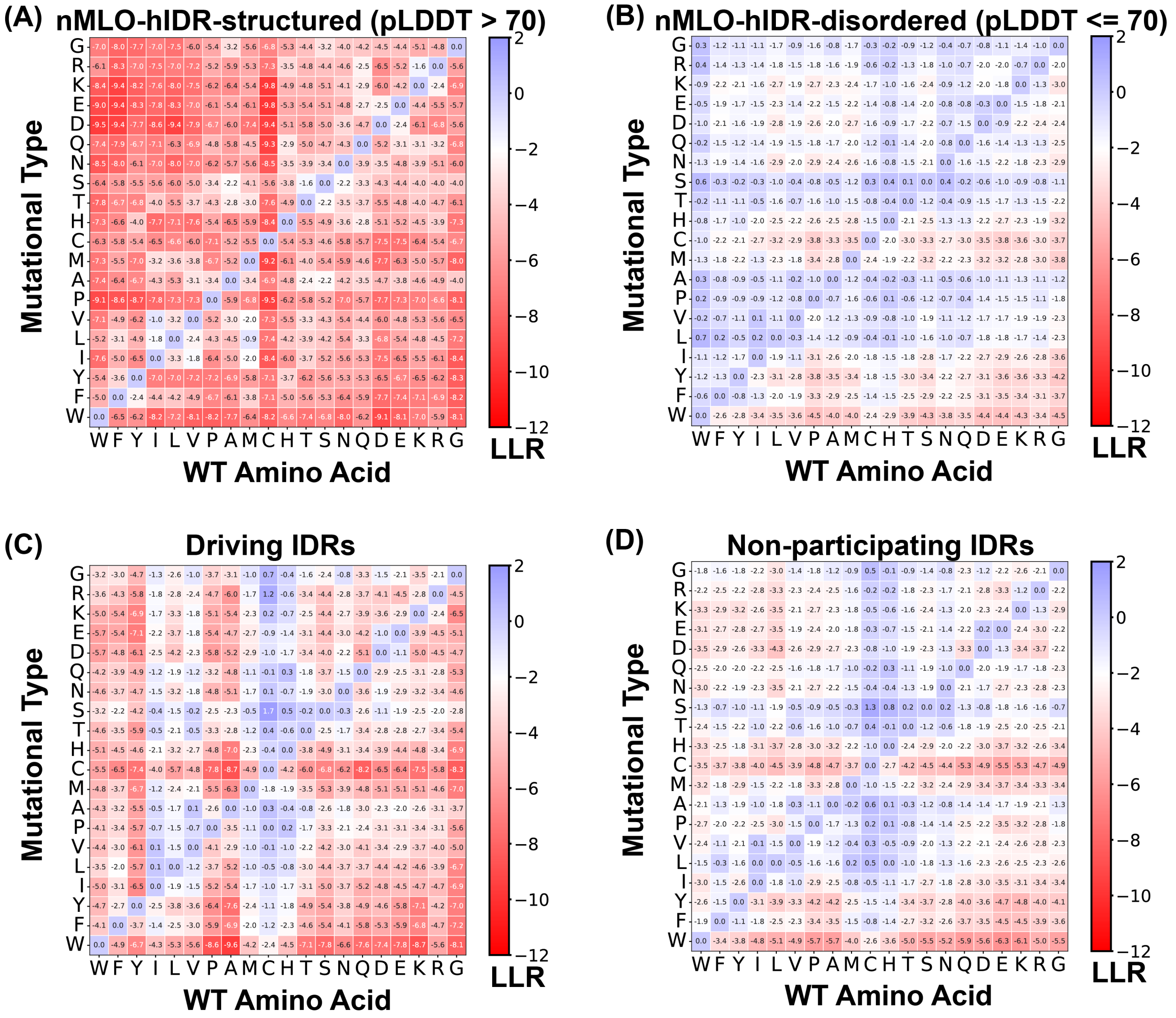


**Figure S9: Mean LLR values for the 20 amino acids, calculated by averaging across all residues of each amino acid type.** The four panels depict averages derived from distinct sets of IDRs: (A) structured residues from nMLO-hIDR ($p$LDDT $> 70$), (B) disordered residues from nMLO-hIDR ($p$LDDT $\leq 70$), (C) driving IDRs from dMLO-IDR, and (D) non-participating IDRs from dMLO-IDR.

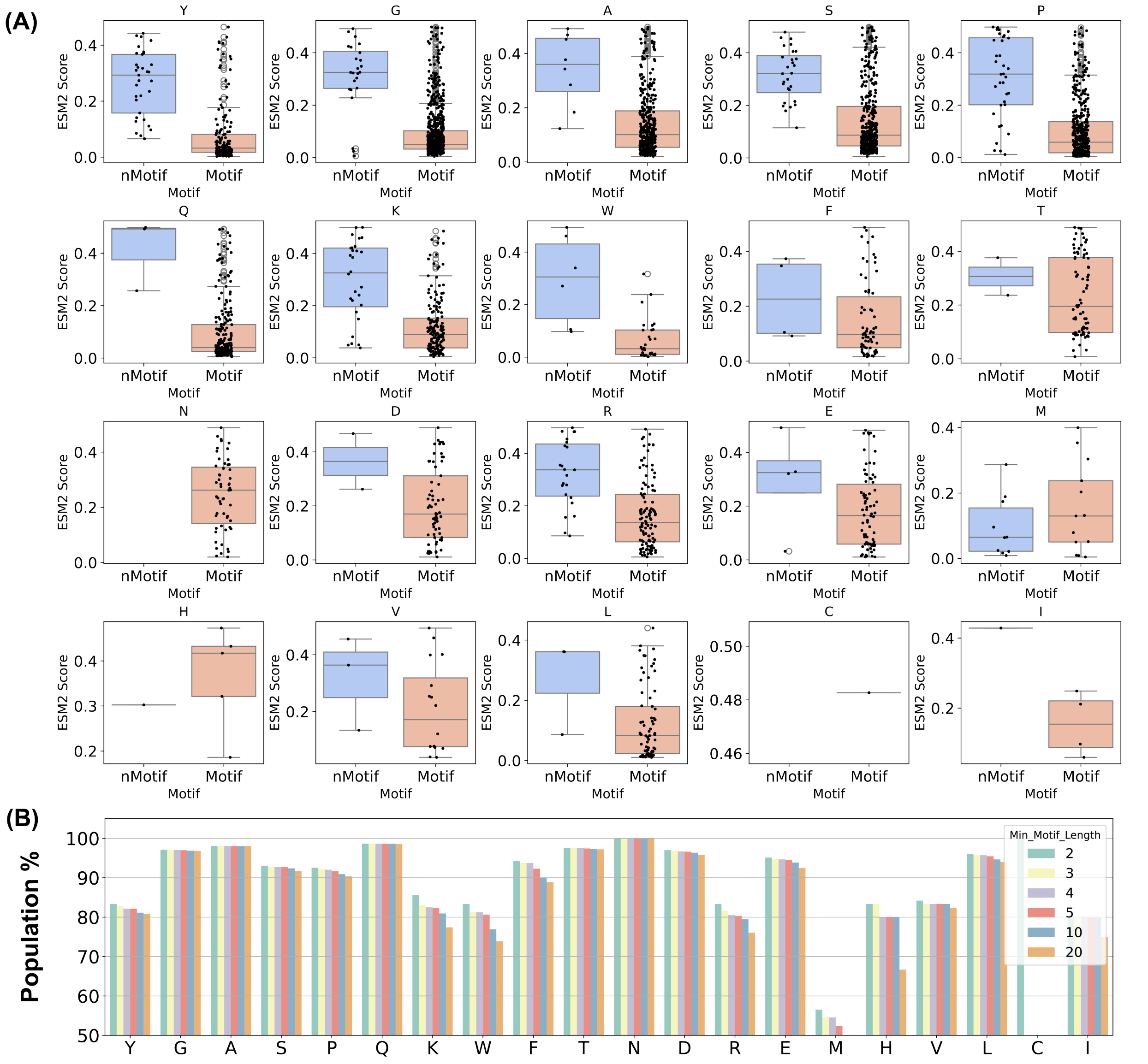


**Figure S10: Distribution of conserved residues in Motifs for driving IDRs.** (A) Box plot illustrating the distribution of conserved amino acids in non-motifs (nMotifs) compared to motifs. Individual data points are displayed as scatter plots. (B) Proportion of conserved amino acids within motifs, evaluated across different minimum motif length thresholds.

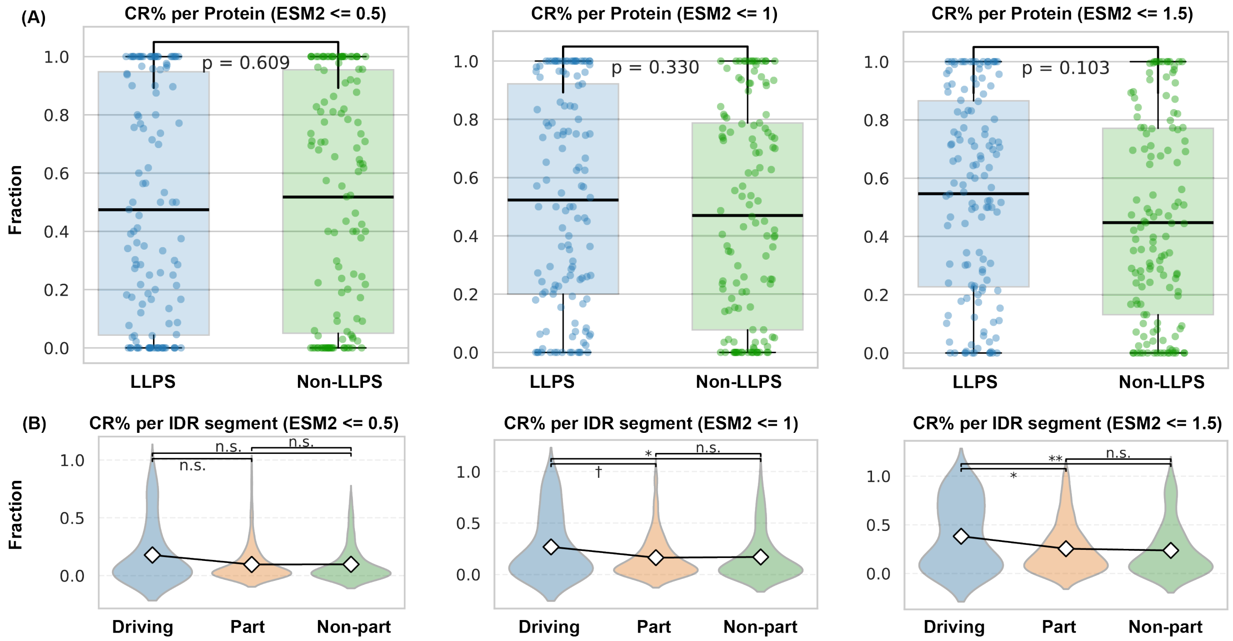


**Figure S11: Statistics of conserved amino acids in regions with varying contributions to phase separation.** (A) Probability of conserved residues occurring in different regions, calculated as the ratio of the number of conserved residues in each region to the total number of conserved residues across the full IDR for each protein. IDRs classified as driving or participating in phase separation (Figure 4B) are grouped under "LLPS," while those not involved are labeled "non-LLPS." The solid black line indicates the mean fraction across all proteins. Conservation was defined using different ESM2 thresholds: $\leq 0.5$, $\leq 1.0$, and $\leq 1.5$, shown in the three panels. (B) Probability of a residue being conserved within each region, computed as the number of conserved residues in a segment divided by the segment's total sequence length. Diamond markers indicate mean values.

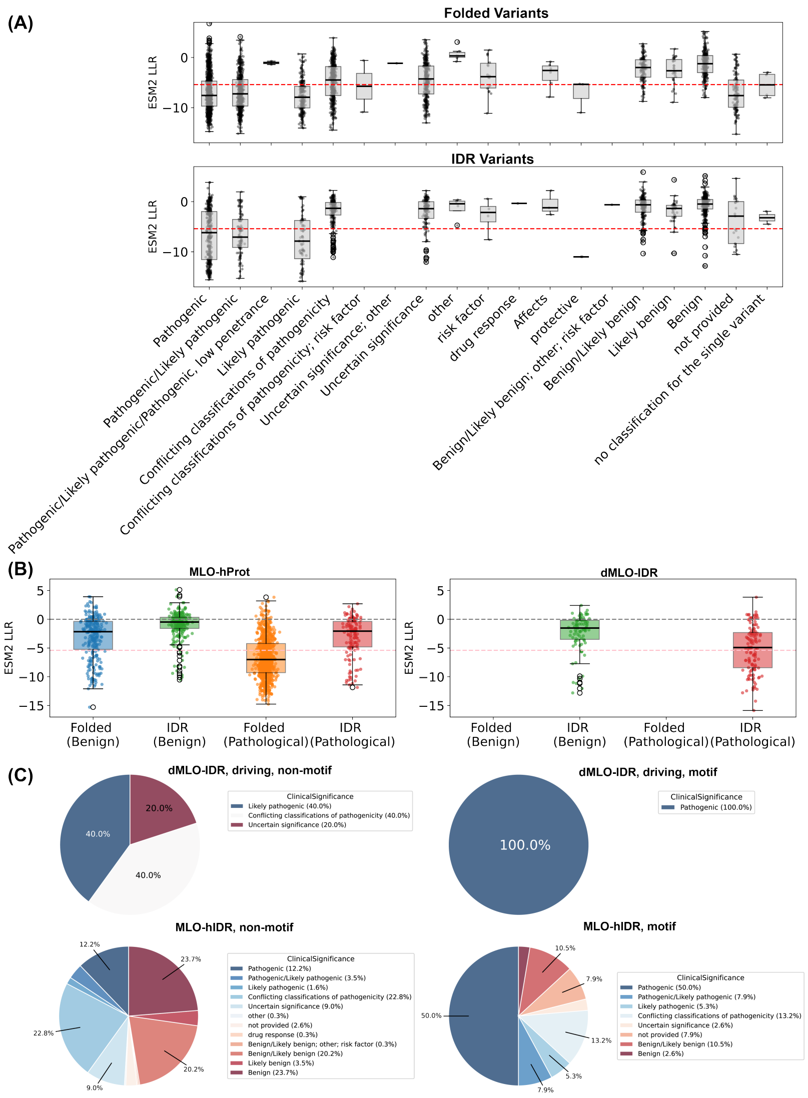


**Figure S12: ESM2 predictions serve as reliable indicators of pathogenic mutations.** Clinical variant data released on March 2024 were used for analysis. (A) Distribution of ESM2 log-likelihood ratio (LLR) values for all single-point mutations in folded (top) and disordered (bottom) residues, stratified by ClinVar clinical significance categories.[S3,S4] Variants are identified in both human proteins and the IDR segments of dMLO-IDR. The red dashed line denotes an LLR value of -5.4, corresponding to an ESM2 Score of 0.5 when uniformly applied across mutations. (B) Comparison of ESM2 LLR distributions across the MLO databases. Variants annotated as "pathogenic/likely pathogenic" are classified as pathogenic, while those labeled "benign/likely benign" are considered benign. The red dashed line again marks the LLR threshold of -5.4. (C) Pie charts illustrating the distribution of ClinVar significance annotations for single-point variants located in motif and non-motif regions of dMLO-IDR and MLO-hProt proteins.

### References


(S1) Orti, F.; Fernández, M. L.; Marino-Buslje, C. MLOsMetaDB, a Meta-Database to Centralize the Information on Liquid–Liquid Phase Separation Proteins and Membraneless Organelles. *Protein Science* **2024**, *33*, e4858.

(S2) Di Domenico, T.; Walsh, I.; Martin, A. J.; Tosatto, S. C. MobiDB: A Comprehensive Database of Intrinsic Protein Disorder Annotations. *Bioinformatics* **2012**, *28*, 2080–2081.

(S3) Landrum, M. J.; Lee, J. M.; Riley, G. R.; Jang, W.; Rubinstein, W. S.; Church, D. M.; Maglott, D. R. ClinVar: Public Archive of Relationships among Sequence Variation and Human Phenotype. *Nucleic Acids Research* **2014**, *42*, D980–D985.

(S4) Landrum, M. J.; Lee, J. M.; Benson, M.; Brown, G. R.; Chao, C.; Chitipiralla, S.; Gu, B.; Hart, J.; Hoffman, D.; Jang, W.; Karapetyan, K.; Katz, K.; Liu, C.; Maddipatla, Z.; Malheiro, A.; McDaniel, K.; Ovetsky, M.; Riley, G.; Zhou, G.; Holmes, J. B.; Kattman, B. L.; Maglott, D. R. ClinVar: Improving Access to Variant Interpretations and Supporting Evidence. *Nucleic Acids Research* **2018**, *46*, D1062–D1067.
